# Dual-site beta transcranial alternating current stimulation during a bimanual coordination task modulates functional connectivity between motor areas

**DOI:** 10.1101/2025.04.04.647211

**Authors:** Mareike A. Gann, Ilenia Paparella, Catharina Zich, Ioana Grigoras, Silvana Huertas-Penen, Sebastian W. Rieger, Axel Thielscher, Andrew Sharott, Charlotte J. Stagg, Bettina C. Schwab

## Abstract

**Background:** Communication within brain networks depends on functional connectivity. One promising approach to modulate such connectivity between cortical areas is dual-site transcranial alternating current stimulation (tACS), which non-invasively applies weak alternating currents to two brain areas.

**Objectives/Hypotheses:** In the current study, we aimed to modulate inter-regional functional connectivity with dual-site tACS to bilateral primary motor cortices (M1s) during bimanual coordination and, in turn, alter behaviour.

**Methods:** Using functional magnetic resonance imaging (fMRI), we recorded participants’ brain responses during a bimanual coordination task in a concurrent tACS-fMRI design. While performing a slow and fast version of the task, participants received one of three types of beta (20 Hz) dual-site tACS over both M1s: in-phase, jittered-phase or sham, in a within-subject, repeated measures design.

**Results:** While we did not observe any significant tACS effects on behaviour, the study revealed a disruptive effect of in-phase tACS on interhemispheric connectivity. Additionally, the two active types of tACS (in-phase and jittered-phase) differed in the task-related M1 connectivity with other motor cortical regions, such as premotor cortex and supplementary motor area. Furthermore, individual E-field strengths were related to functional connectivity in the in-phase condition.

**Conclusions:** Dual-site beta tACS over both M1s altered functional connectivity between motor areas. However, this effect did not translate to the behavioural level, possibly due to compensatory mechanisms.

**Highlights:**

1. Interhemispheric connectivity between the primary motor cortices was targeted with dual-site beta tACS during bimanual coordination.
2. In-phase as compared to sham tACS disrupted interhemispheric connectivity during task performance.
3. In-phase and jittered-phase tACS led to different task-related functional connectivity patterns within the motor network.
4. The relationship between individual in-phase E-field strengths and interhemispheric connectivity depended on task demand.
5. There were no significant tACS effects on behavioural performance of bimanual coordination, possibly due to compensatory mechanisms.

**Graphical abstract:** 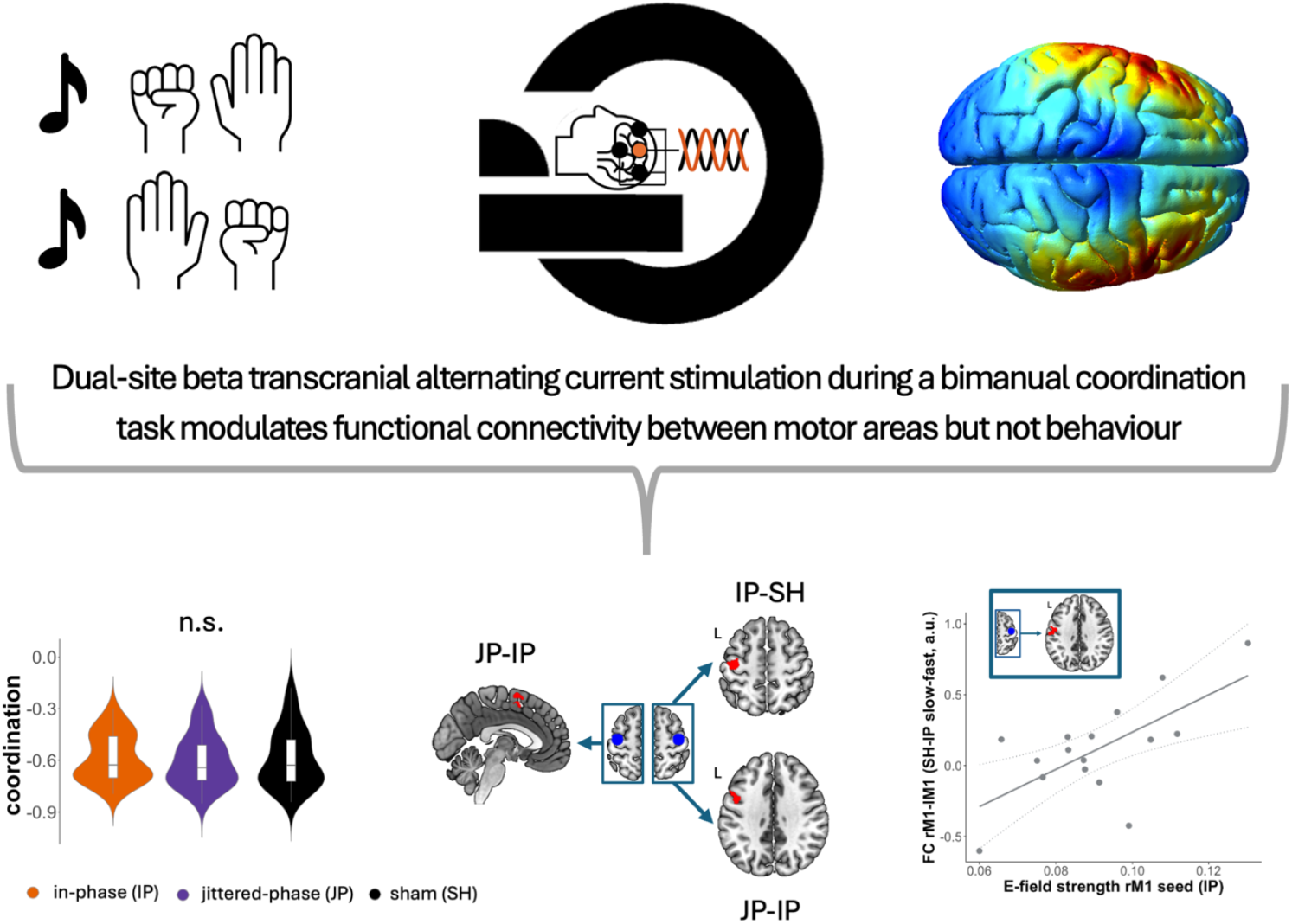

## 1 Introduction

Communication within networks of anatomically distinct brain regions is crucial for supporting human cognition across the breadth of behaviours (1,2). Communication within these networks depends on structural and functional connections between different brain regions in the network. Functional connectivity (FC) can be shaped on a short time scale and therefore provides an essential addition to structural connectivity to support relatively rapid neuroplastic changes. For example, FC in motor networks has been linked to bimanual coordination (3–7), with stronger interhemispheric coupling - specifically in the beta band - relating to better behavioural performance (3).

Yet, although many studies have investigated the correlation between FC and behaviour across a range of functional domains, the degree to which FC between brain regions *causally* underpins the correlated behaviour is still unclear. To determine causality, interventions which specifically modulate FC are required. One promising approach to modulate FC between cortical areas is transcranial alternating current stimulation (tACS), a form of non-invasive brain stimulation (NIBS), that applies weak alternating currents to the scalp to entrain neural responses (8–11).

A recent tACS study (12) showed that beta (20Hz) tACS over the left primary motor cortex (M1) modulated the pattern of FC within the motor network, as observed with functional magnetic resonance imaging (fMRI) during rest. However, multiple questions remain to understand how tACS is able to modulate motor FC. Firstly, it is likely that tACS interacts with networks differently depending on how engaged they are in performing a task (so-called state-dependency, (13–15)). To simultaneously observe brain responses and apply tACS during a motor task, fMRI is the method of choice, given that, compared to electroencephalography (EEG) or magnetoencephalography (MEG) there are no problematic tACS-induced artefacts (16). Nevertheless, very few studies have directly investigated the effects of tACS on active motor networks with fMRI. Secondly, applying tACS at two different sites opens the possibility to compare the different phase shifts of the applied currents and to potentially selectively modulate FC between the two sites. However, tACS has not often been used to modulate brain responses in more than one node within the motor network. Finally, tACS is typically used at an intensity of 2mA peak-to-peak or lower (17). tACS applied at higher current amplitudes is thought to entrain ongoing neural oscillation towards the externally applied frequencies more effectively than lower currents (17–19), potentially offering a more robust method to modulate FC.

Here, we aimed at investigating the role of communication in the beta band in inter-regional FC within the motor network. We did this in the context of a bimanual coordination task that requires meaningful communication between the two M1s. Bimanual coordination (i.e., the interplay of both hands) relies on interhemispheric FC within the beta band (13-30Hz) between several motor areas (3,4,6). Indeed, motor beta interhemispheric FC can be disrupted in healthy aging as well as in neurological conditions such as stroke or Parkinson’s disease (20–23) and modulating beta functional connectivity is therefore a highly promising target for rehabilitative therapies. Thus, we wanted to determine whether tACS interacts with a network differentially depending on how active that network is. To do this, we used a bimanual task requiring two levels of engagement, elicited by slow and fast movements.

Specifically, we used high-definition dual-site, in-phase, 20Hz tACS over both M1s at a 4mA peak-to-peak amplitude, which is thought to entrain FC between both homologous areas to the stimulation frequency and consequently increase FC. This, in turn, should lead to improved behavioural performance. We also investigated a dual-site tACS protocol with a changing phase-relationship between both hemispheres (jittered-phase tACS) at the same intensity, which is thought to lead to similar sensations but different modulations of interhemispheric FC (24). In the context of bimanual coordination, we expected such jittered-phase beta dual-site tACS to disrupt FC and consequently decrease task performance.

Altogether, in this fMRI study we employed focalised tACS on the two most important nodes of the active motor network to modulate and read out inter-regional FC as well as the associated bimanual coordination behaviour at varying task demands.

## 2 Methods

### 2.1 Participants

All volunteers provided written informed consent to participate in accordance with the Central University Research Ethics Committee approval (University of Oxford; CUREC ethics reference R68812). All participants were eligible for tACS and MRI, reported no current psychiatric or neurological disorders, and had normal or corrected-to-normal vision. Eighteen young healthy right-handed participants completed both sessions of the current study (age 24.11±3.28 years, range 20-31 years; 10 females). One participant only completed two of the three task runs due to technical difficulties. Additionally, six volunteers withdrew during the study due to timing constraints, non-compliance with instructions, or feeling uncomfortable during tACS application and were not included in any analysis. One volunteer was subsequently identified as an outlier in behaviour (>3 standard deviations above the coordination values of the rest of the sample) and was therefore excluded. Therefore, the analyses included data from 17 or 16 volunteers. All participants received financial compensation for their participation.

### 2.2 Experimental design

Volunteers participated in two experimental visits (at least one week apart; 15.61±9.53 days between the two visits). During the first visit, participants were familiarised with tACS (as applied with two round rubber electrodes of 40mm diameter on EEG locations F4 and PO4) and the bimanual coordination task. They also completed two runs of the bimanual coordination task to ensure that they understood the task demands.

During the second visit, the volunteers were familiarised with the tACS setup as used inside the MRI scanner and practiced the bimanual coordination task again outside the MRI scanner. Inside the MR scanner, volunteers performed three task runs (of 16 task blocks each) with concurrent tACS of approximately 13 mins each. Task scans were interleaved with T1- and T2-weighted structural scans (Figure 1A). After each concurrent tACS-task run, volunteers were asked about the strength of the sensations that they felt due to the brain stimulation from 1=absent to 5=strong. After the MRI scans, volunteers filled in an extensive questionnaire on sensations during tACS (see Supplementary Figure S1 and Supplementary Results).

**Figure 1.**
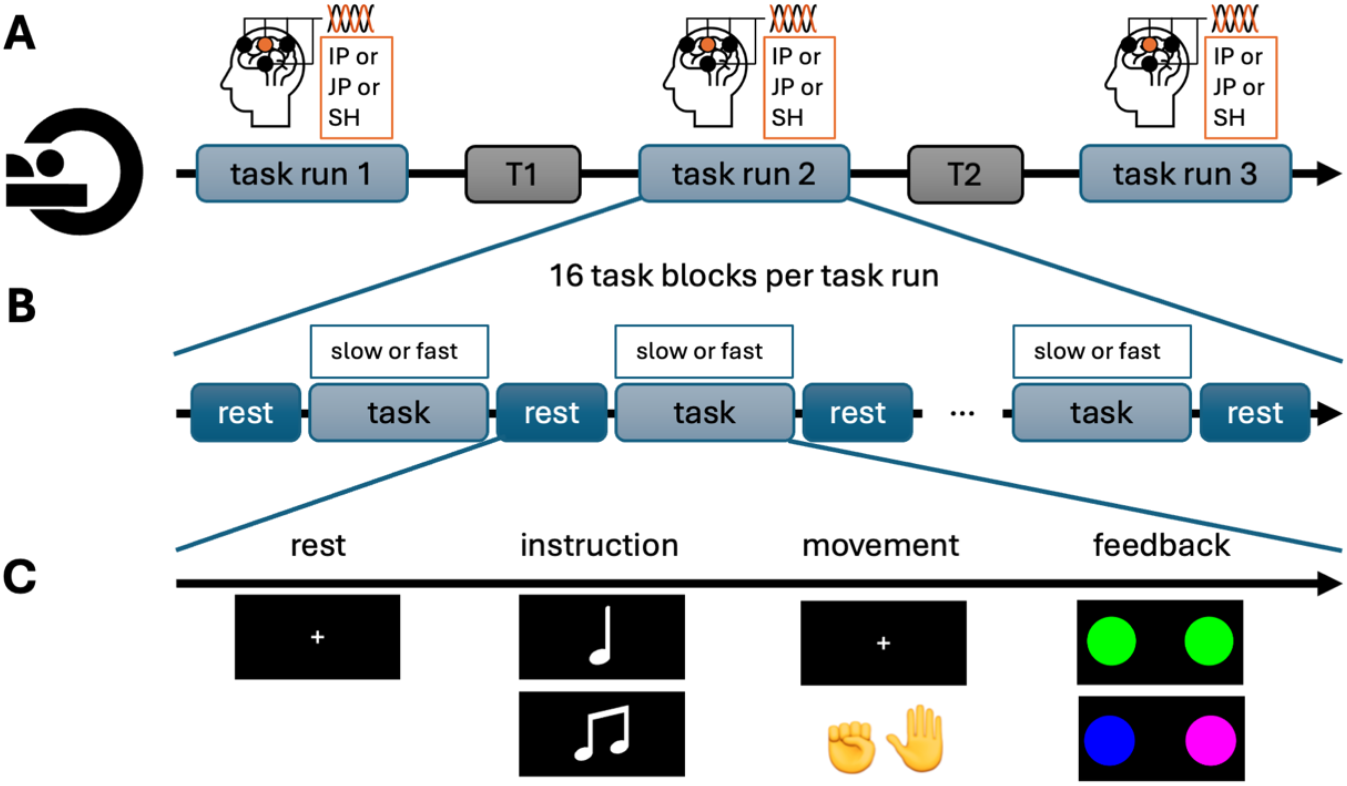
Overview of the MR and task setup. (A) The MR part of the study during visit 2 consisted of three task runs with concurrent tACS and two structural scans (T1 and T2 for T1- and T2-weighted data, respectively). (B) Each task run consisted of 16 task blocks, alternating between slow and fast task blocks, interleaved with rest blocks. (C) Each task block following a rest block started with a 2s instruction screen, then the volunteers performed the alternating movements in sync with the presented metronome sound for 30s and saw feedback on the applied force for 1s. MR – magnetic resonance, tACS – transcranial alternating current stimulation, IP – in-phase tACS, JP – jittered-phase tACS, SH – sham tACS.

### 2.3 Bimanual coordination task

Participants performed asymmetrical alternating movements using MR-compatible grip force sensors (Grip Force Bimanual Fiber Optic Response Pad, HHSC-2×1-GRFC-V2, Current Designs Inc., Philadelphia, PA, United States) synchronized to metronome sounds, presented at 3.8Hz for the slow and 6Hz for the fast task blocks. Participants were required to make smooth, well-coordinated alternating hand movements, with each hand squeezing the force grip sensors in sync with a metronome sound. The task was implemented in MATLAB R2016b (The MathWorks Inc., Natick, MA, United States) using PsychToolbox (25).

The required force was individually adjusted. Therefore, in each session, participants squeezed the force grip sensors as hard as possible to calculate their maximum voluntary force (MVF) for each hand. For the task runs participants were instructed to use 7-13% of their MVF. Each task run consisted of 16 alternating fast and slow task blocks, with the starting order counterbalanced across the group (Figure 1B). Blocks began with a 2s instruction indicating the speed (fast/slow), followed by a 30s task period during which participants performed coordinated movements in sync with the metronome sounds while fixating on a central cross. Participants then received 1s feedback on the force used (green for correct (7-13% MVF), pink for excessive force (>13% MVF) and blue for insufficient force (<7% MVF); Figure 1C). The rest blocks, where a fixation cross was shown for 15s, preceded each task block.

Task performance was quantified using Pearson’s correlation between the force-time course of both hands (see Supplementary Figure S2 for an exemplary time course of one task block). During the first visit and during task training of the second visit, experimenters gave feedback until this correlation was sufficient (defined as r<-0.53 in the slow and r<-0.37 in the fast conditions; these threshold values were determined as 20% above the mean values observed in a behavioural pilot study; Pearson’s r of −1 would represent perfectly alternating movements of the left and right hands). Feedback was also given to ensure that the speed of movement was 1.4 - 2.4Hz in slow blocks (target 1.9Hz) and 2.5 - 3.5Hz in the fast blocks (target 3Hz).

Linear mixed-effects models with speed and stimulation conditions as fixed-effect factors and volunteers as random-effect factors were performed in R (R Core Team, Vienna, Austria) using the lme4 package (26), with the mean Pearson correlation for each task speed and stimulation condition for each participant as the dependent variable. Any additional covariates were individually added as fixed effect factors to the model.

### 2.4 Transcranial alternating current stimulation

Visit 1 employed a simpler two-electrode montage at F4 and PO4. Two 40mm diameter electrodes were positioned using Ten20 paste. Visit 2 employed an 8-electrode montage, with 40mm diameter electrodes at C3/4, F3/4 and T7/8 and 23mm diameter electrodes at P3/4 (see Figure 2A). The current at the central electrodes (i.e. C3/4) was phase-shifted 180° from the corresponding outer electrodes in both 3-in-1 montages, allowing us to target the M1s with low current leakage across hemispheres. Impedance at each electrode was kept below 10kΩ.

**Figure 2.**
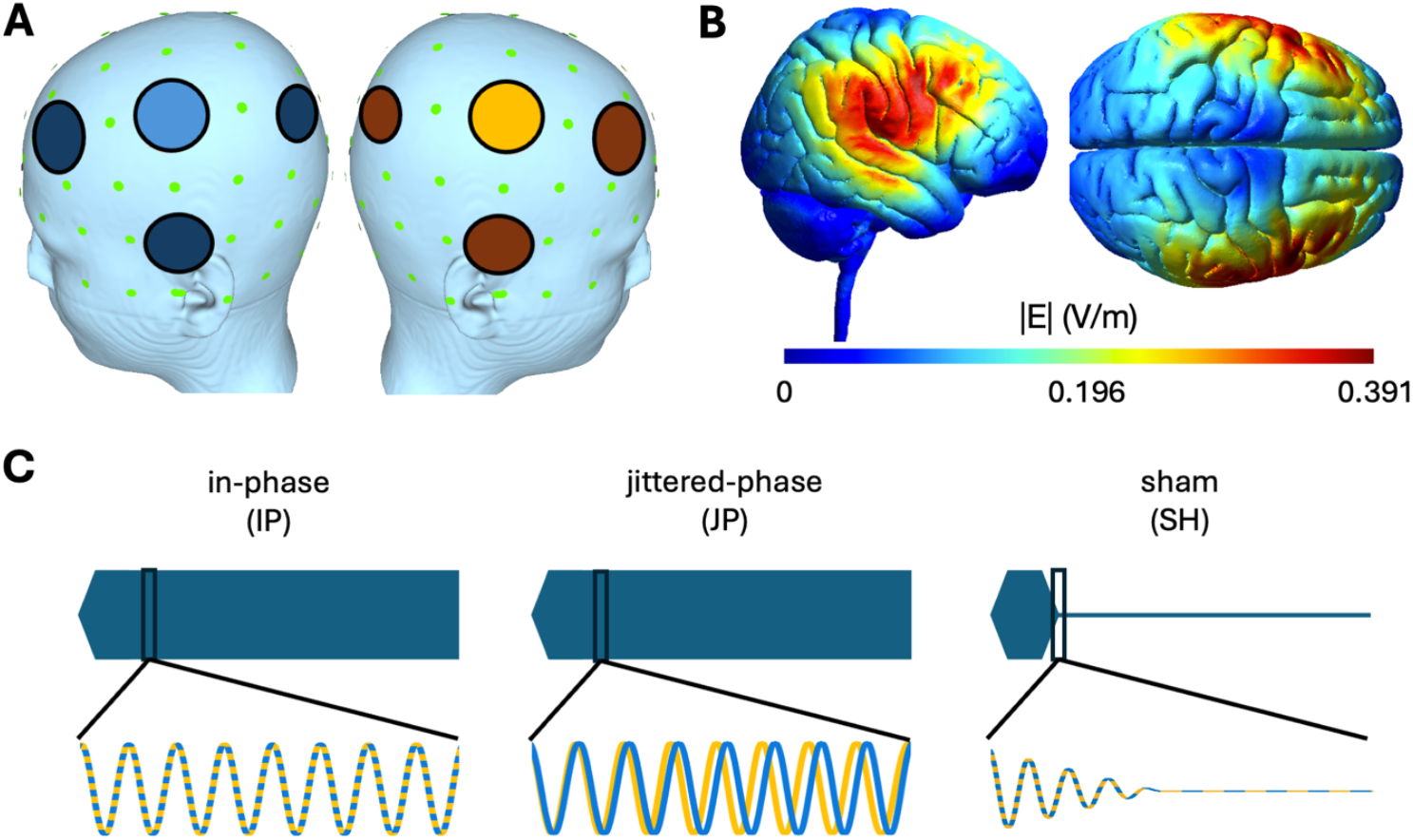
Transcranial alternating current stimulation (tACS) setup. (A) tACS montage shown for the left and right hemisphere during the tACS-fMRI setup. Colours represent the phase shift: the inner electrodes over C4 (light orange) and over C3 (light blue) are shifted by 180° as compared to the respective outer electrodes, depicted in darker colours. This montage led to an E-field distribution on the MNI template for in-phase (IP) stimulation at 4mA peak-to-peak as shown in (B). (C) Schematic overview of the tACS conditions: The waveforms of inner electrodes of the tACS, i.e. C3 as shown in blue and C4 as shown in orange, follow the same 20Hz sinusoidal currents for the in-phase (IP) condition and two currents that independently changed their frequency between 19.5 and 20.5Hz for the jittered-phase (JP) condition. The sham (SH) conditions consisted of a short (10s) IP stimulation followed by a 3s ramp-down to zero. All conditions start with a 3s ramp-in period.

In visit 1, the tACS current was increased in a stepwise fashion, to ensure that participants could tolerate 4mA peak-to-peak comfortably. In visit 2, we performed a short (60s) familiarisation tACS stimulation outside the MR scanner. During the MR-task runs, tACS was applied at 4mA peak-to-peak at the C3/4 electrodes, with a 3s ramp-up and ramp-down.

Three stimulation conditions were tested: in-phase (IP), jittered-phase (JP) and sham (SH) tACS. IP stimulation used 20Hz sinusoidal currents with phase offset zero between the two hemispheres (see Figure 2B for simulated E-field distributions). For JP stimulation two currents were applied that independently changed their frequency between 19.5 and 20.5Hz, leading to continuous interhemispheric phase-shifts (Figure 2C). For SH stimulation, IP stimulation was applied for 10s followed by a 3s ramp-down. The order of stimulation conditions was counterbalanced across the group, and participants were blinded to the stimulation conditions.

#### 2.4.1 E-field simulations

To quantify the E-field in M1 and to investigate whether interindividual differences influenced behaviour and fMRI readouts we conducted E-field simulations using SimNIBS 4.0.1 (27) on both the MNI template and individual anatomical scans. Individual head models were constructed from T1 and T2 scans using Charm (28). tACS electrode artefacts were removed manually using ITK-SNAP (http://www.itksnap.org, (29)). The skin-air boundaries of the segmentations were then smoothed using a Gaussian smoothing kernel with a radius of 3mm prior to running the simulations. We extracted the maximum E-field strength and the absolute normal component across the entire grey matter (GM) sheet and within the M1s (region 4 from the HCP-MMP1 atlas (30)) from both the individual and MNI surface simulations. We also extracted the mean E-field within the GM in the seeds of left and right M1 as used for the psychophysiological interaction (PPI) analysis (see below), as well as in the precentral gyrus (as taken from the AAL atlas (31)).

### 2.5 Magnetic resonance imaging

#### 2.5.1 MR acquisition

Whole-brain MR images were acquired on a Siemens Prisma 3T MRI system equipped with a 64-channel head coil. Task-related functional MR images were acquired using a multi-band echo planar imaging (MB-EPI, (32–34)) sequence (766 volumes, TR=1030ms, TE=37.4ms, MB acceleration factor 6, flip angle=60°, 72 coronal slices, FOV=208×208×144mm^3^, voxel size=2×2×2mm^3^). Additionally, one field map was acquired for every participant (TR=607ms, TE1=4.92ms, TE2=7.38, flip angle=46°, 62 coronal slices, FOV=210×210×155mm^3^, voxel size=2.5×2.5×2.5mm^3^). High-resolution T1-weighted 3D structural images (TR=1900ms, TE=3.96ms, flip angle: 8°, FOV=232×256×192mm^3^, voxel size=1×1×1mm^3^, 192 transversal slices) were acquired with a Magnetisation Prepared Rapid Acquisition Gradient Echo sequence and T2-weighted structural images (TR=2500ms, TE=275ms, flip angle: 8°, FOV=272×272×191mm^3^, voxel size=0.8×0.8×0.8mm^3^, 224 sagittal slices) without fat suppression.

#### 2.5.2 Task-related MR analyses: activation

FMRI analyses were performed using FMRIB Software Library (FSL, Oxford; (35)). The following preprocessing steps were applied: B0 field map unwarping, motion-correction with MCFLIRT (36), slice-time correction, brain extraction using BET, grand-mean intensity normalisation of the entire 4D dataset by a single multiplicative factor, and high-pass temporal filtering with a 100s cut-off. The T1 images were bias-field corrected using fsl_anat. FMRI images were linearly registered to the individual T1 scan, via a task-run specific reference volume, and then non-linearly to the MNI152 template using FNIRT (35). FIX (FMRIB’s ICA-based Xnoiseifier (37)) was applied to denoise the data. Following denoising, the fMRI data was smoothed with a 5mm FWHM Gaussian kernel and then modelled with a general linear model (GLM) with 5 explanatory variables (EVs): (EV 1) slow task blocks, (EV 2) fast task blocks, (EV 3) instruction screen, (EV 4) feedback screen and (EV 5) any instances of force values above 5% of the individual maximum force during rest blocks. These EVs were convolved with the double-gamma haemodynamic response function, temporal derivatives were added, and temporal filtering was applied. Standard and extended motion parameters were added as confounds. The different sessions per participant were combined in a second-level FEAT analysis. Detailed descriptions of the contrasts are provided in the Supplementary Information.

We first identified a mean task activation network, independent of tACS and speed condition, using Mixed Effects (FLAME 1+2 (38)) with cluster-forming thresholds of Z = 2.3 and P = 0.01 on the group level. To define the M1 seeds for subsequent FC analyses, we then applied a pre-threshold mask of the bilateral A4ul regions (upper limb motor regions in the precentral gyrus) from the Brainnetome atlas (https://atlas.brainnetome.org, (39)) with cluster-forming thresholds of Z = 2.3 and P = 0.01.

Next, we compared the speed conditions (slow-fast) independent of tACS, using Mixed Effects (FLAME 1+2 (38)) with cluster-forming thresholds of Z = 2.3 and P = 0.01 on the group level. To contrast the different tACS and speed conditions, we ran a second-level analysis for each participant separately with fixed effects. At the group-level, we then conducted a Mixed Effects analysis using FLAME 1+2 (38). To investigate the effects of tACS specifically on the motor-related cortex, we used a pre-threshold mask including motor cortical areas (bilateral pre- and postcentral areas, bilateral supplementary motor areas as defined in the AAL atlas (31)), with cluster-forming thresholds of Z = 2.3 and P = 0.01.

#### 2.5.3 Task-related MR analyses: functional connectivity

Psychophysiological interaction (PPI) analyses were performed on the denoised and smoothed data to examine tACS-related changes in FC of task-specific peaks in bilateral M1s. To do this, we defined the left and right M1 seeds 8mm radius spheres around the peak coordinates from the activation analysis detailed above.

Time courses were extracted from denoised data for M1 seeds and control regions in the cerebrospinal fluid (CSF) and white matter (WM). The first-level GLM for the FC analyses across speed conditions (slow+fast) included nine EVs: (EV 1) the psychological factor (slow and fast task blocks), (EV 2) the physiological factor (seed time course), (EV 3) the interaction term of the psychological and physiological factor (PPI), (EV 4) the instruction screen, (EV 5) the feedback screen and (EV 6) the force used during rest blocks (see above), the time course of the (EV 7) left CSF and the (EV 8) right CSF and of the (EV 9) WM. For the FC analyses comparing speed conditions (slow-fast) the psychological factor (EV 1) consisted of the slow task blocks modelled with +1 and the fast task blocks modelled with −1 and an additional EV (EV 10) was added for all task blocks (slow+fast). The EVs for the psychological factor, instruction screen, feedback screen, rest block force and those including task blocks were convolved with the double-gamma haemodynamic response function, temporal derivatives were added and temporal filtering applied. Additionally, standard and extended motion parameters were added as confounds. Second-level and group-level analyses were conducted as detailed above for the activation analyses.

#### 2.5.4 Task-related MR analyses: regression analyses

Group-level analyses incorporated covariates, including sensation rating differences between conditions as covariate of no interest as a follow-up on the tACS effect on interhemispheric FC (see Results section). Additionally, we added mean E-field values within PPI seed regions during IP as a covariate of interest in a more restrictive mask, including only the contralateral precentral and postcentral gyrus (AAL atlas). As JP stimulation has comparable E-field strengths to IP stimulation (24), we assumed the same values for JP. Furthermore, even though no E-fields were applied during SH, we formally assigned the same E-field strengths for SH as simulated for IP. We expected that we would not find any relationship between the E-field magnitude and the FC for the SH condition. Non-normally distributed covariates underwent Box-Cox transformation. All covariates were demeaned before inclusion in fMRI analyses.

## 3 Results

### 3.1 E-field simulations quantified M1 targeting

We first wanted to know whether our tACS montage was able to target the M1s bilaterally. Individual surface simulations revealed maximum field strengths of 0.45±0.047V/m across the grey matter, and 0.432±0.050V/m in M1s (99.9th percentile). The mean E-field values in the PPI M1 seeds were 0.095±0.016V/m and 0.089±0.017V/m for left and right seeds respectively.

### 3.2 20Hz tACS to bilateral M1 did not modulate behaviour

Next, we investigated whether bi-hemispheric 20Hz tACS modulated behaviour on our bimanual coordination task. A linear mixed-effects analysis of bimanual coordination, with one factor of speed (fast, slow) and one factor of stimulation (IP, JP, SH) revealed a significant main effect of speed (X^2^(1)=20.594, *p*<0.001), such that participants performed better during the slow blocks compared to the fast ones (t(86.1)= −4.746, *p*<0.0001, Figure 3A), as would be expected. However, there was no main effect of stimulation (X^2^(2)=1.0232, *p*=0.60) and no speed by stimulation interaction (X^2^(2)=1.8575, *p*=0.40).

**Figure 3.**
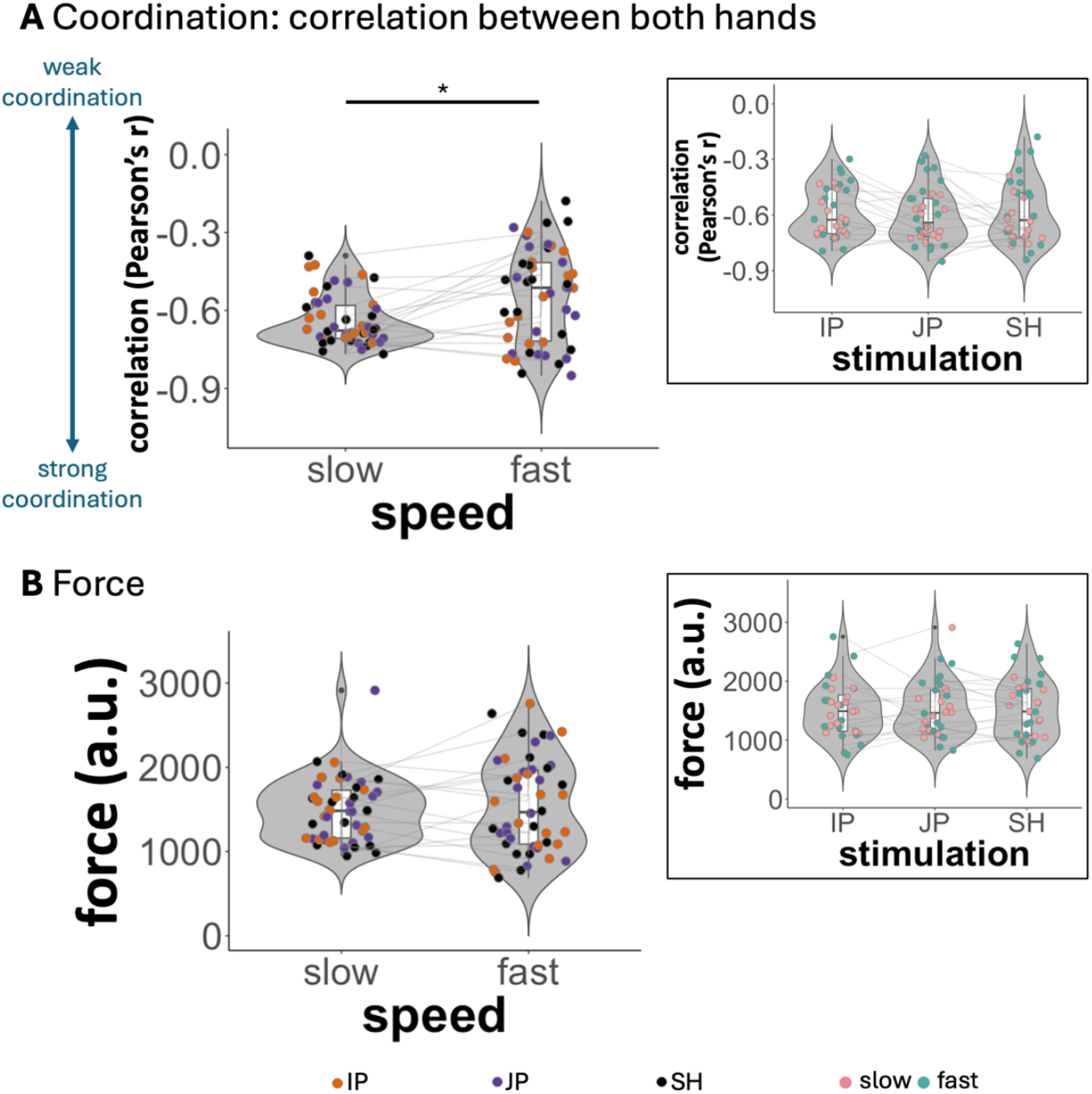
Statistical model results for coordination and force. (A) For coordination, we observed a significant speed effect, showing better coordination (i.e., a correlation value (Pearson’s r) of −1 would represent perfectly alternating movements of the left and right hands) during the slower as compared to the faster task blocks, but no stimulation-related effects. (B) The force levels were neither influenced by speed or stimulation. Filled circles (jittered horizontally within the respective violin plot) represent individual data and lines connect these across conditions for each participant. The shape of the violin plots and the included boxplots depict the distribution of the data. Asterisks (*) indicate significant statistical tests (*p*<0.05). a.u. – arbitrary unit, tACS – transcranial alternating current stimulation, IP – in-phase tACS, JP – jittered-phase tACS, SH – sham tACS

To determine whether the amplitude of the applied E-field was associated with behavioural effects, we added a covariate of interest as a fixed effect to our models. There was no main effect of the E-field in the precentral gyrus (X^2^(1)= 3.1165, *p*=0.08), of the E-field in the PPI seeds (i.e., adding the E-fields of the left and right PPI seeds) (X^2^(1)= 0.204, *p*=0.65), nor of the sensation ratings (X^2^(1)= 0.0381, *p*=0.85) on the coordination of the two hands.

To investigate whether tACS led to differences in force used in the task, we performed a linear mixed-effects analyses which revealed no speed by stimulation interaction (X^2^(2)= 0.3575, *p*=0.84), no main effect of stimulation (X^2^(2)= 0.0448, *p*= 0.98) and no main effect of speed (X^2^(1)= 1.2671, *p*= 0.26; see Figure 3B).

### 3.3 tACS did not modulate task-related fMRI activation

We then wanted to determine whether bi-hemispheric 20Hz tACS modulated task-related neural activation. We first determined which brain regions were active during task performance, independent of tACS condition or task level (see Supplementary Table S1 and Supplementary Figure S3). As would be expected in this type of task, we saw task-related activations in multiple cortical and subcortical motor areas, including the primary motor cortex, premotor cortex, supplementary motor area (SMA), striatum and cerebellum. Next, we determined the brain regions that differed between task levels, independent of tACS conditions (see Supplementary Table S1 and Supplementary Figure S4). We observed higher activation levels in the slow condition compared to the fast one, in multiple cortical and subcortical areas, including bilateral precentral and postcentral regions, superior parietal regions, SMA, caudate, thalamus, putamen, central opercular cortex, pallidum, insula and cerebellum. We observed higher activation in fast compared to slow conditions in several areas, including the frontal regions, (para)cingulate regions, occipital cortex, precuneus, inferior temporal areas, (para-)hippocampus, amygdala, insula and left cerebellum.

Critically, there were no significant differences in neural activation patterns in response to the motor task among the three stimulation conditions.

### 3.4 General task-related fMRI functional connectivity of the M1 seeds

To address our hypothesis that 20Hz tACS to bilateral M1s modulates interhemispheric FC, we performed a PPI analysis. As detailed in the Methods section, we created 8mm radius spherical seed regions in the left and right M1s, centred on as the mean centre of gravity of the task-related activity in each M1 separately (see Supplementary Table S2 and Figure 4A).

**Figure 4.**
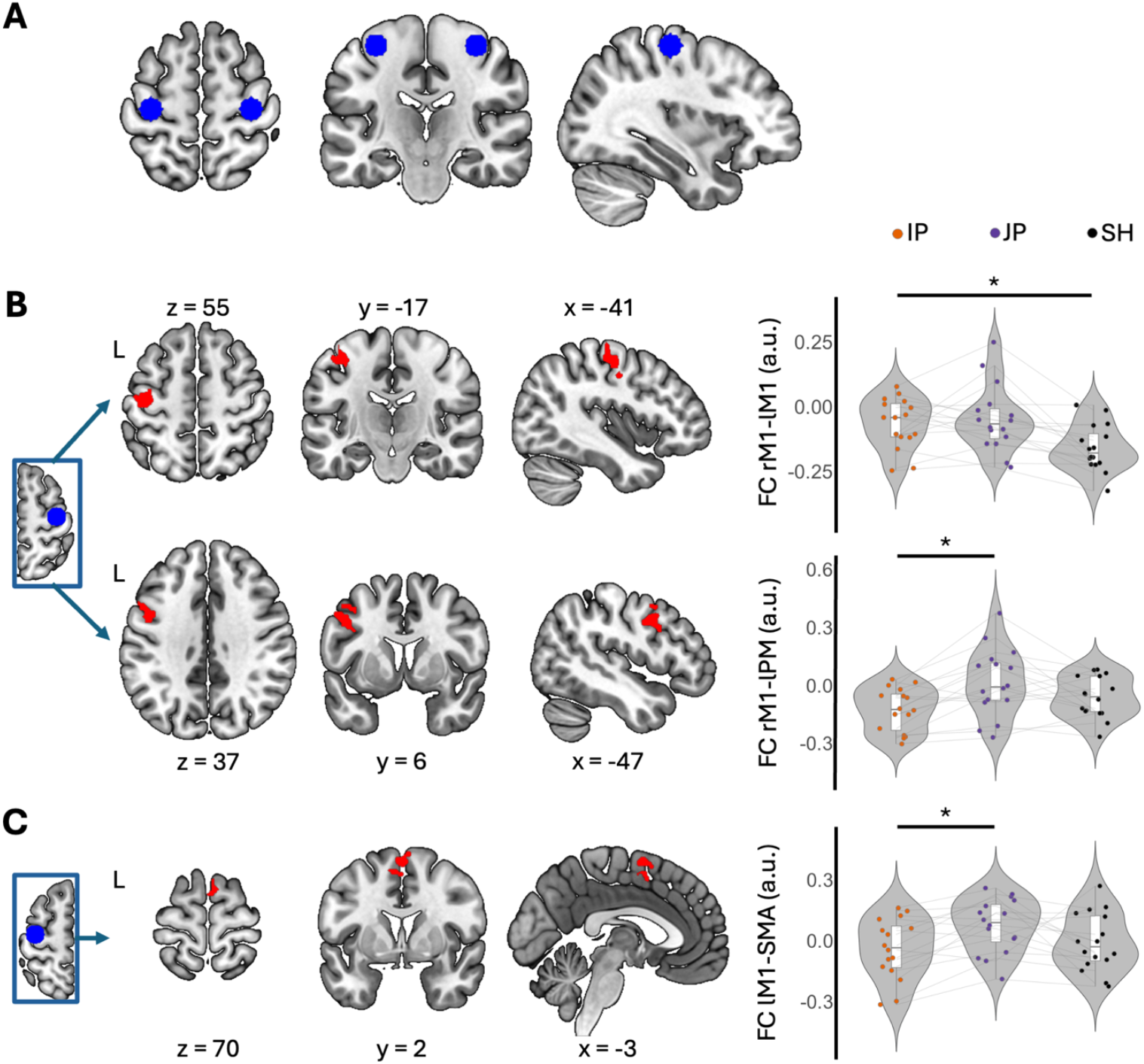
Effects of tACS on functional connectivity. (A) Primary motor cortex (M1) seeds for the fMRI functional connectivity (FC) analyses: The 8mm radius spheres used for the FC analyses are depicted in blue. (B) FC seeded in the right M1 (rM1) differed significantly in a left M1 (lM1) cluster between in-phase (IP) and sham (SH) conditions. The FC in SH was lower, i.e. negative, than in IP. Additionally, the FC seeded in the right M1 differed significantly in a left premotor (lPM) cluster between in-phase (IP) and jittered-phase (JP) conditions, with higher FC in JP. (C) FC seeded in the left M1 differed significantly in a cluster in the supplementary motor area (SMA) between IP and JP conditions, with higher FC in JP. The spheres and clusters are displayed on a T1-weighted template image (neurological convention). Filled circles (jittered horizontally within the respective violin plot) represent individual data and lines connect these across conditions for each participant. The shape of the violin plots and the boxplots depict the distribution of the data. Asterisks (*) indicate the contrast from which the fMRI clusters resulted. fMRI – functional magnetic resonance imaging

First, we investigated the FC of the two motor cortices, independent of tACS condition or speed. Directly correlating the PPI time-courses revealed a significant negative FC between left and right M1 (see Supplementary Table S3.1 and S3.3 as well as Supplementary Figure S5A and B). This negative FC pattern was stronger (i.e., had a larger absolute value) in the slow compared to the fast blocks (see Table S3.2 and S3.4 as well as Supplementary Figure S5C and D). Additionally, both M1s were significantly negatively correlated with extensive regions stretching through the M1s and postcentral gyri bilaterally, and significantly positively correlated with clusters within ipsilateral M1. For the right M1 seed, the latter, positively correlated, ipsilateral M1 cluster extended into the SMA. Additionally, the FC between left M1 and SMA was higher in the fast compared to the slow blocks (see Supplementary Table S3 and Supplementary Figure S5).

### 3.5 tACS modulated task-related functional connectivity between M1s

We next investigated what effect bilateral 20Hz tACS would have on M1 FC patterns. Comparing FC patterns across speed conditions (i.e. combining slow and fast blocks) between different tACS conditions demonstrated that FC between the right and left M1 seeds was significantly different in the sham and IP conditions, such that there was significantly less negative FC between left and right M1s in IP compared with sham tACS (see Figure 4B upper part and Table 1.1). There was no significant difference between JP and IP conditions, nor between JP and SH conditions at the chosen thresholds.

**Table 1.**
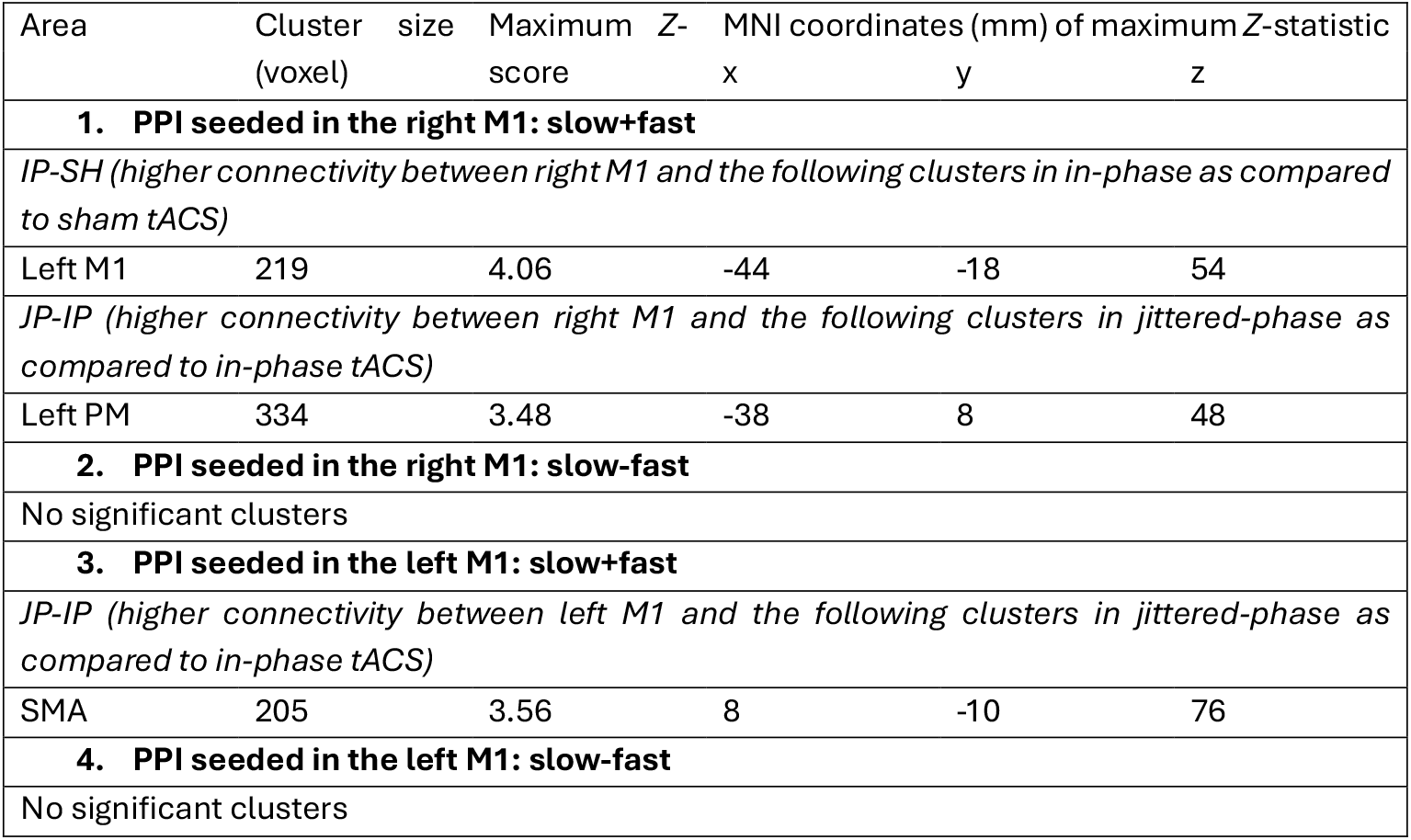
Functional connectivity comparing tACS and speed conditions. Clusters were determined with cluster-forming thresholds of Z = 2.3 and P = 0.01 within the cortical motor regions (precentral and postcentral gyrus, premotor cortices (PM), supplementary motor area (SMA)). MNI – Montreal Neurological Institute.

We observed a difference between sham and active stimulation conditions in terms of the sensations evoked (see Supplementary Information). To ensure that the differences in FC described above could not be simply explained by differences in sensation, we ran a control analysis including sensation rating as a covariate of no interest. The significant cluster described above remained significant after controlling for sensation ratings (see Table 2.1 in comparison to Table 1.1).

**Table 2.** Functional connectivity comparing tACS and speed conditions with sensations used as covariate of no interest: i.e., the corresponding difference between the sensation ratings of the pair of conditions used in the fMRI contrast. Clusters were determined with cluster-forming thresholds of Z = 2.3 and P = 0.01 within the cortical motor regions (precentral and postcentral gyrus, premotor cortices (PM), supplementary motor area (SMA)). MNI – Montreal Neurological Institute.

| Area | Cluster size (voxel) | Maximum Z-score | Z-x | MNI coordinates (mm) of maximum Z-statistic y | z |
| --- | --- | --- | --- | --- | --- |
| **1. PPI seeded in the right M1: slow+fast**<br>**covariate: difference in sensation ratings** |  |  |  |  |  |
| *IP-SH, covariate of no interest: sensations IP-SH* |  |  |  |  |  |
| Left M1 | 199 | 3.98 | -44 | -18 | 54 |
| *JP-IP, covariate of no interest: sensations JP-IP* |  |  |  |  |  |
| Left PM | 311 | 3.43 | -38 | 8 | 48 |
| **2. PPI seeded in the left M1: slow+fast**<br>**covariate: difference in sensation ratings** |  |  |  |  |  |

| *JP-IP, covariate of no interest: sensations JP-IP* |  |  |  |  |  |
| --- | --- | --- | --- | --- | --- |
| SMA | 305 | 3.73 | 6 | -10 | 74 |
| Left postcentral gyrus | 218 | 3.98 | 38 | -38 | 62 |

Next, we wanted to know whether the effects of tACS on task-related FC were dependent on the level of engagement of the network in the task being performed. We therefore used the speed condition difference (slow-fast) as the psychological factor of the PPI analyses described previously. For either the right or left M1 seed PPIs, we observed no significant clusters when comparing the different tACS conditions between speed conditions (slow-fast).

### 3.6 tACS modulated task-related functional connectivity between M1s and the rest of the motor network

We then wanted to investigate whether bilateral M1 tACS led to a change in how the M1 connected with the rest of the motor network. Key to our hypothesis was that there would be a different effect of IP and JP on FC. Indeed, IP and JP differentially modulated FC between right M1 and left premotor cortex (PM), such that the FC was stronger during JP than during IP (see Table 1.1 and Figure 4B lower part). The same analysis seeded in the left M1 revealed a difference in FC between left M1 and the supplementary motor area (SMA) when comparing JP and IP conditions (see Table 1.3 and Figure 4C). Here, again, FC between left M1 and SMA was stronger during JP than during IP. The tACS effects in FC did not differ between the speed conditions for either the right or left M1 seeds.

**Table 3.**
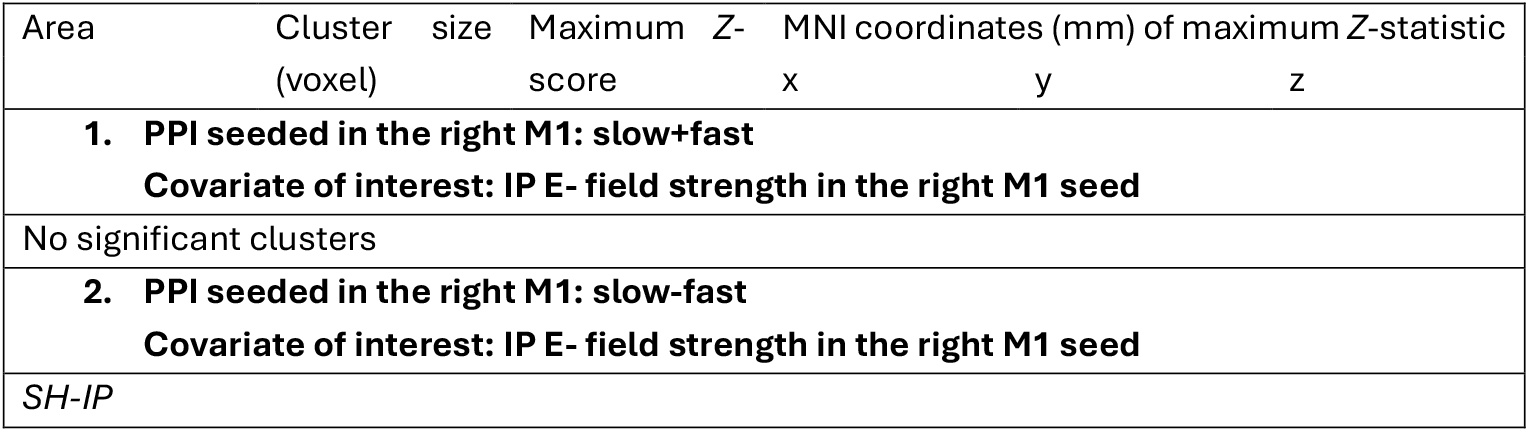
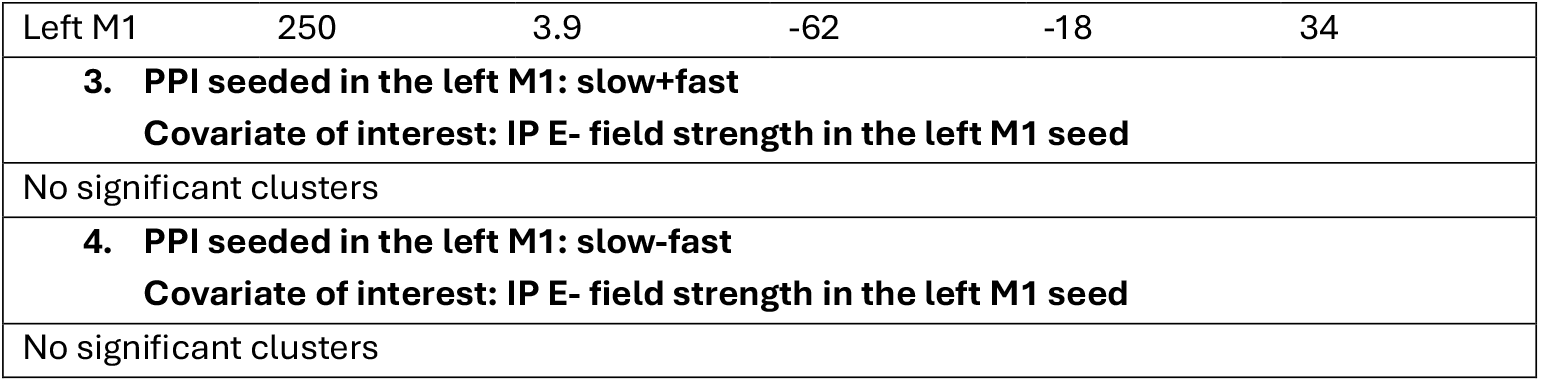
Functional connectivity comparing tACS and speed conditions with IP E-field in the PPI seed used as covariate. Clusters were determined with cluster-forming thresholds of Z = 2.3 and P = 0.01 within the contralateral primary sensorimotor regions (precentral and postcentral gyrus). MNI – Montreal Neurological Institute.

The significant clusters described above remained significant after controlling for sensation differences evoked by tACS (see Table 2 in comparison to Table 1). In addition, we observed an additional cluster in the left post-central gyrus where intrahemispheric FC with the left M1 PPI seed significantly differed between IP and JP tACS (see Table 2.2).

### 3.7 Relationship between functional connectivity and individual E-field strengths

Finally, we tested whether there was a direct relation between E-field strength and interhemispheric FC changes due to tACS. We therefore added the individual simulated E-field strength for IP within the GM of the seed region as a covariate of interest to the PPI analyses. When considering the task demand, i.e. comparing the slow and fast conditions (PPI slow-fast), we observed a significant difference between IP and SH conditions in the correlation between the applied IP E-field magnitude in the right M1 and right M1 FC with the contralateral left M1 (see Table 3.2 and Figure 5A). Interestingly, this effect was driven by the significant relationship between IP E-field in the right M1 seed and the FC of the right M1 seed with the left M1 cluster in the IP slow condition (SH slow: t_(14)_=1.116, *p*_*uncorrected*_=0.2832; SH fast: t_(14)_=-0.93509, *p*_*uncorrected*_=0.3656; IP slow: t_(14)_=-3.1706, *p*_*uncorrected*_=0.006807; IP fast: t_(14)_=0.92527, *p*_*uncorrected*_=0.3705; see Figure 5B). Specifically, higher IP E-field values related to stronger negative FC during the IP slow condition.

**Figure 5.**
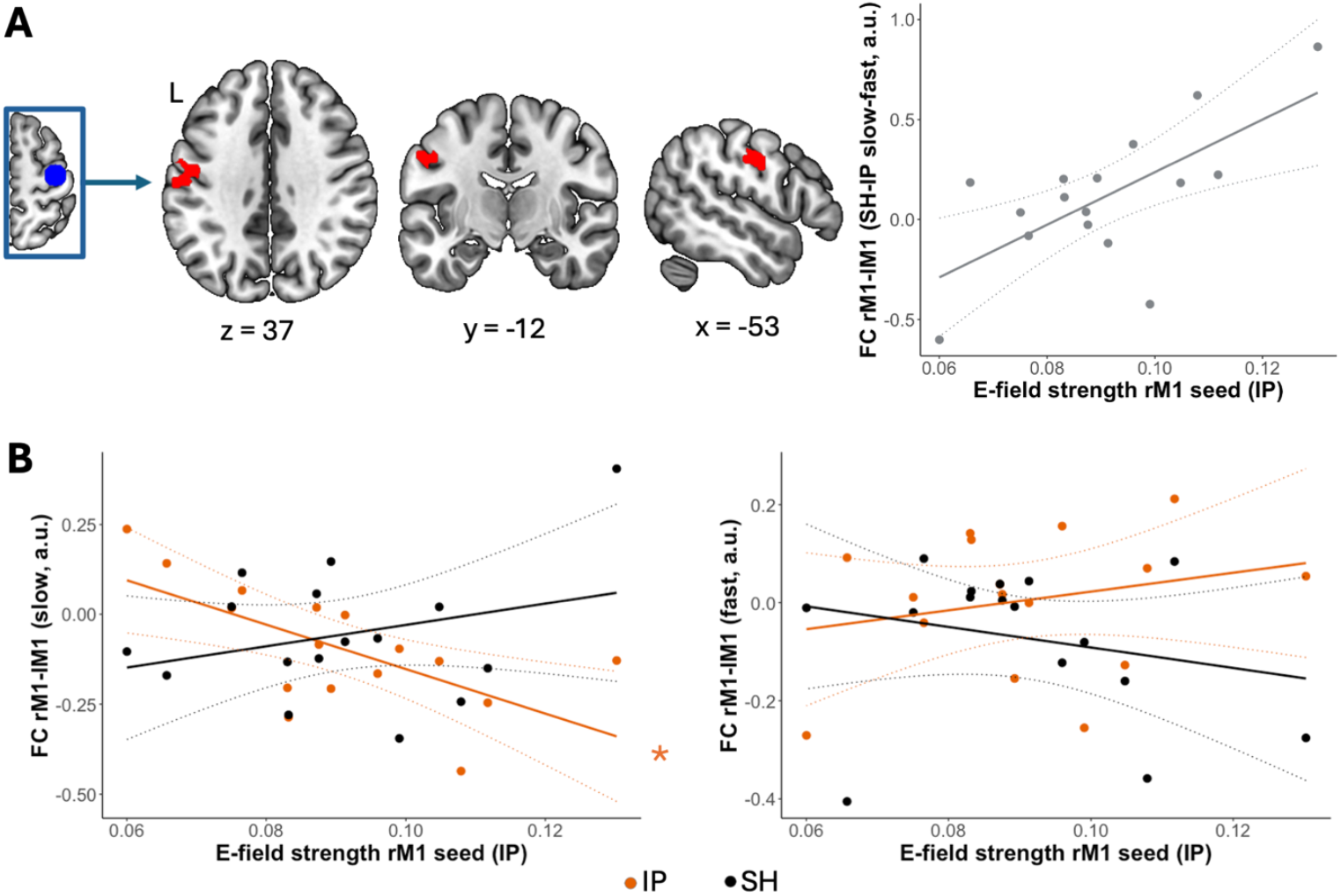
Modulation of relationship between IP E-field strength in right M1seed and functional connectivity by tACS. (A) The relationship between the applied IP E-field magnitude in the right M1 (rM1) and right M1 functional connectivity (FC) with the contralateral left M1 (lM1) differed significantly between IP and SH conditions when comparing the slow and fast conditions. The clusters are displayed on a T1-weighted template image (neurological convention). (B) Follow-up tests showed a significant relationship between IP E-field strength and FC only in the IP slow condition. Asterisks (*) indicate significant relationships (Pearson’s correlation with *p*<0.05) between the FC and covariate values in individual conditions (e.g. IP slow). Note that the raw E-field strength values are depicted on the plots rather than the demeaned values used as covariates for the analyses. Filled circles represent individual data, solid lines depict the linear regression fits, and dotted lines represent the 95% prediction intervals of the linear function. M1 – primary motor cortex, tACS – transcranial alternating current stimulation, IP – in-phase tACS, SH – sham tACS, a.u. – arbitrary unit

## 4 Discussion

In this study, we combined a high-definition dual-site transcranial alternating current (tACS) setup at 4mA with a concurrent functional magnetic resonance imaging (fMRI) design, to investigate changes in functional connectivity (FC) during tACS. We targeted the bilateral primary motor cortices (M1s) during a bimanual motor coordination task, thus enabling the investigation of active motor networks. Contrary to our expectation, we did not observe any significant tACS effects on behaviour. However, we found a disruptive effect of in-phase (IP) tACS on interhemispheric FC. Additionally, the task-related M1 FC with other motor cortical regions (premotor cortex (PM) and supplementary motor area (SMA)) differed between the two active types of stimulation. Finally, individual E-field strengths correlated with interhemispheric FC in the IP condition, depending on task demand.

### 4.1 Disrupted interhemispheric connectivity

Our results suggest that IP tACS disrupts interhemispheric FC between M1s compared to sham stimulation. This deviates from our hypotheses expecting an increase in interhemispheric FC between the M1s during IP tACS and a decrease in FC during jittered-phase (JP) tACS. As the underlying FC between both M1s was negative during the execution of this specific asymmetrical coordination task, the application of IP tACS might have disrupted negative FC towards positive FC by applying the identical stimulation above both regions. Similarly, unifocal beta tACS disrupted the relationship between interhemispheric M1 FC and the rest of the motor network (12). A recent study reported increased interhemispheric inhibition between the M1s (as measured with transcranial magnetic stimulation) after IP 20Hz tACS (40). While it is difficult to directly translate this result to fMRI FC, this might point to a more complex effect of IP tACS on interhemispheric FC, which is worth exploring with multimodal approaches. Future studies should also investigate the effects of anti-phase tACS, which might be, speculatively, more closely related to negative FC as measured with fMRI.

Alternatively, the disruptive effect of IP tACS might be explained by our use of a fixed tACS frequency (20Hz) for all the participants. Instead of entraining the underlying oscillations to the applied 20Hz rhythm, we might have introduced a strong and consistent noise factor disrupting the underlying negative FC between both M1s. Thus, individualized IP tACS might enhance ongoing FC and associated behaviour. Similarly, previous studies showed that individualized-frequency alpha tACS over both M1s increased resting-state connectivity (8), while fixed-frequency unifocal beta tACS disrupted the relationship between interhemispheric M1 FC and the rest of the motor network (12). In addition, a recent study has shown that the interplay of interregional conduction delays and stimulation frequency is critical for the effects of dual-site tACS on stimulation-outlasting EEG FC changes (41). Heterogeneity in the individual conduction delays between the M1s could thus further contribute to variability in our data.

Conversely, JP tACS did not disrupt interhemispheric FC at the chosen thresholds, as compared to sham or JP tACS. While JP tACS might also introduce noise and result in numerically less strong negative interhemispheric FC (as seen in Figure 5B), this response may not be consistent enough to lead to significant effects. In line with this, a recent study showed that IP tACS resulted in stronger effects on FC than JP tACS in a resting-state tACS-EEG study (24).

### 4.2 Potentially compensatory effects in response to tACS

Modifying brain dynamics may come along with compensatory and adaptive effects. Accordingly, we found further tACS-related responses that were revealed in the brain imaging analyses. Specifically, the FC between M1 seeds and PM as well as SMA differed between the two active tACS conditions. While these effects do not directly explain how the disrupted interhemispheric FC as compared to the sham condition was counteracted in the IP condition, they may point to broader compensatory mechanisms involving different recruitment strategies within the motor cortical network under different types of dual-site tACS to maintain behavioural performance levels. The spread of tACS effects through the (cortical and subcortical) motor networks is in line with previous work in M1 tACS-fMRI studies using beta (8,12) as well as mu (42) tACS. Additionally, PM and SMA might be more strongly coupled to M1 in JP than in IP condition due to the lower predictability of the JP stimulation. The SMA is thought to be involved in internally-guided movements, reflecting the coordination between both hands in our task, and the PM in externally-guided movements, reflecting the speed-adjustment towards the metronome sounds (e.g. (43–45)). The increased connectivity between M1 and these higher-order processing regions may therefore reflect a compensatory mechanism to maintain behaviour in the presence of a higher cognitive load required to counteract the unpredictable noise introduced by JP tACS.

Interestingly, tACS effects on FC did not differ between task demand levels, suggesting that initial differences in recruitment across fast and slow conditions did not lead to speed-specific tACS modulations. Importantly, even though the participants rated the sensations stronger for active than for sham conditions, the observed effects were not statistically driven by these sensations. However, to avoid possible influences of different sensations in different tACS conditions, future studies could focus on better blinding mechanisms, such as applying a numbing cream underneath the tACS electrodes.

### 4.3 Relationship between interhemispheric connectivity and the E-field strength

The effects of tACS are known to be variable, at least in part due to the varying intensity of the electrical fields within the cortex. Therefore, we investigated whether the variation between participants in the E-fields applied to M1 may have influenced patterns of FC changes. We demonstrated speed-specific tACS modulation of the relationship between interhemispheric FC and IP E-field strength. Specifically, there was only a significant relationship between IP E-field strength and interhemispheric FC during IP tACS in the slow condition. Our analyses demonstrated that the slow condition led to greater activity in the motor cortex and showed stronger negative FC, than the fast condition. These findings might seem counterintuitive but may result from more focus on “proper” motor control in the slow condition and from less control, decreased coordination and a greater cognitive load in the fast condition.

The strong and consistent interhemispheric bilateral M1 recruitment during slow coordination might then be more susceptible to disruption than that seen in the fast condition, as there is more room for such disruptive modulations. Specifically, lower E-fields in the IP condition showed stronger disruption of the negative FC in the lateral left M1. This finding suggests that, in this specific region, we do not see a disruptive effect of IP tACS overall, but possibly a disruptive effect emerges only at lower E-fields. This hypothesis might be supported by recent theories on the working mechanism of tACS, suggesting that tACS at lower E-fields desynchronises the underlying neural responses rather than synchronising responses to the applied current, as would be expected for higher E-fields (18). Such desynchronisation might promote disruptive effects of the underlying FC. While JP tACS produced similar E-fields as IP tACS, the JP stimulation pattern might not show similar effects on the (de)synchronisation of underlying neural responses.

### 4.4 Asymmetry in tACS effects

Notably, while the bimanual coordination task execution as well as the stimulation were symmetrical, the results reveal an asymmetry in tACS effects. Specifically, tACS-related FC patterns differed between left and right M1 seeds. This may be explained by the right-handedness of our participants. Additionally, motor skills do not recruit motor regions symmetrically due to hemispheric dominance (46). Similarly, a current M1 tACS-fMRI study also showed asymmetrical results for mu tACS effects on task-related FC seeded in the targeted left and right M1 (42).

### 4.5 Active tACS did not modulate motor performance or activation

Neither IP nor JP tACS modulated motor performance during the bimanual coordination task. These results suggest that dual-site tACS over both M1s does not necessarily lead to changes in motor performance. One possible explanation is that bimanual coordination is not causally linked to FC of the M1s. A second option is that the observed modulations in FC were not strong enough to alter behaviour in the investigated task setup. However, a potentially more plausible explanation, especially considering the disrupted interhemispheric FC in the IP condition and further tACS effects on FC when comparing the active tACS conditions, is the use of compensatory mechanisms and differential recruitment of these other brain regions. Furthermore, while the tACS current is stronger here than in most other studies, the frequency was not individualized. Even though we used high-definition montages, E-fields were not only focused in the M1s, and showed inter-participant variability. All these considerations might explain a high variability in effects leading to a null result on the behavioural level. Similarly, the effects of bilateral M1 beta tACS showed inconsistent results in the literature: while some researchers observed behavioural effects on motor tasks using bilateral beta (40,47) M1 tACS, others could not (48–50). At this point the literature on bilateral M1 tACS in the beta band during motor tasks is stills scarce, especially for high-definition montages. More research is needed to pinpoint the exact working mechanisms to reveal dual-site M1 beta tACS effects on motor tasks and whether these differ across different types of tasks (such as bimanual motor control, visuomotor tasks or motor learning) and phase-lags between the hemispheres. Additionally, other frequency bands, such as alpha and mu have shown promise to modulate motor behaviour with bilateral M1 tACS (42,47).

Similarly, while we observed tACS effects on FC, we did not see any tACS-related effects on fMRI activation. Presumably tACS, and specifically dual-site tACS, alters FC more than activation as observed with fMRI. This is in accordance with previous literature on tACS-fMRI using unifocal beta tACS over M1 showing tACS modulations of FC in the absence of effects on activation (12). While alpha tACS-fMRI studies showed changes in BOLD activation, these effects were subtle and might depend strongly on the specific target regions, phase-lags between dual-site tACS regions and appear to be state-dependent (51,52). Furthermore, a recent study (53) indicated that tACS may have the most dominant effect on long-range projections rather than on local spiking. While this new finding is generally in line with our results of tACS modulating connectivity more than local activity, it could also imply that projections to other regions may have been stimulated additionally.

## 5 Conclusions

Our study showed that high-definition dual-site beta-tACS over both primary motor cortices altered functional connectivity between motor areas. However, this effect did not translate to the behavioural level, possibly due to compensatory mechanisms. Consequently, dual-site tACS has the general potential for connectivity modulation, which is very important for neuroscientific studies as well as for clinical translations, not limited to the motor system. At the same time, more research is necessary to understand the complex mechanisms of dual-site tACS.

## Author contributions

**Mareike A. Gann**: Conceptualization; Resources; Data curation; Software; Formal analysis; Investigation; Visualization; Methodology; Writing - original draft; Writing - review and editing **Ilenia Paparella**: Investigation; Methodology; Writing - review and editing **Catharina Zich**: Conceptualization; Resources; Writing - review and editing **Ioana Grigoras**: Methodology; Data curation; Software; Formal analysis; Writing - review and editing **Silvana Huertas-Penen:** Data curation; Software; Writing - review and editing **Sebastian W. Rieger:** Conceptualization; Methodology; Writing - review and editing **Axel Thielscher:** Software; Writing - review and editing **Andrew Sharott:** Conceptualization; Funding acquisition; Writing - review and editing **Charlotte J. Stagg**: Conceptualization; Resources; Data curation; Software; Formal analysis; Supervision; Funding acquisition; Validation; Investigation; Project administration; Writing - original draft; Writing - review and editing **Bettina C. Schwab**: Conceptualization; Resources; Data curation; Software; Formal analysis; Supervision; Funding acquisition; Validation; Investigation; Project administration; Writing - original draft; Writing - review and editing

## Data Availability

At the time of publication, group fMRI statistical outputs mapped to standard space, as well as image and data processing pipelines from our study will be made freely available at the Data Sharing Platform of MRC Brain Network Dynamics Unit (https://data.mrc.ox.ac.uk/).

## Acknowledgements

This work was supported by the German Research Foundation (DFG, grant number SCHW 2023/2-1 to B.C.S.), the Medical Sciences Internal Fund: Pump-Priming (grant number 0011905 to A.S.) of the University of Oxford, the Medical Research Council UK (MC_UU_00003/6 to A.S.), a Wellcome Trust Senior Research Fellowship (224430/Z/21/Z to C.J.S), the Sofina Boël Fund for Education and Talent (to I.P.), the F.R.S.-FNRS (to I.P.), the National Institute for Health Research (NIHR) Oxford Biomedical Research Centre and the NIHR Oxford Health Biomedical Research Centre (NIHR203316). AT was supported by the Lundbeck Foundation (grants R313-2019-622 and R244-2017-196) and the German Research Foundation (Research Unit 5429/1 (467143400), TH 1330/6-1 and TH 1330/7-1). The views expressed are those of the author(s) and not necessarily those of the NIHR or the Department of Health and Social Care. The Wellcome Centre for Integrative Neuroimaging is supported by core funding from the Wellcome Trust (203139/Z/16/Z and 203139/A/16/Z). For the purpose of Open Access, the author has applied a CC BY public copyright licence to any Author Accepted Manuscript (AAM) version arising from this submission. Additionally, we acknowledge the receipt of software for using multiband fMRI sequences from the University of Minnesota Center for Magnetic Resonance Research.

We thank all involved students and colleagues for assistance with data collection. We thank the WIN radiographers for imaging assistance. We thank Saad Jbabdi and Eik Vettorazzi for analysis assistance.

## References

1. Mišić B, Sporns, Olaf. From regions to connections and networks: new bridges between brain and behavior. Curr Opin Neurobiol. 2016;

2. Perich MG, Rajan K. Rethinking brain-wide interactions through multi-region ‘network of networks’ models. Curr Opin Neurobiol. 2020 Dec;65:146–51.

3. Serrien DJ, Brown P. The functional role of interhemispheric synchronization in the control of bimanual timing tasks. 2002;

4. Grefkes C, Eickhoff SB, Nowak DA, Dafotakis M, Fink GR. Dynamic intra- and interhemispheric interactions during unilateral and bilateral hand movements assessed with fMRI and DCM. NeuroImage. 2008 Jul;41(4):1382–94.

5. Daffertshofer A, Peper C (Lieke) E, Beek PJ. Stabilization of bimanual coordination due to active interhemispheric inhibition: a dynamical account. Biol Cybern. 2005 Feb;92(2):101–9.

6. Donchin O, Gribova A, Steinberg O, Bergman H, Vaadia E. Primary motor cortex is involved in bimanual coordination. Nature. 1998 Sep;395(6699):274–8.

7. Houweling S, Van Dijk BW, Beek PJ, Daffertshofer A. Cortico-spinal synchronization reflects changes in performance when learning a complex bimanual task. NeuroImage. 2010 Feb;49(4):3269– 75.

8. Bächinger M, Zerbi V, Moisa M, Polania R, Liu Q, Mantini D, et al. Concurrent tACS-fMRI Reveals Causal Influence of Power Synchronized Neural Activity on Resting State fMRI Connectivity. J Neurosci. 2017 May 3;37(18):4766 LP – 4777.

9. Bergmann TO, Karabanov A, Hartwigsen G, Thielscher A, Siebner HR. Combining non-invasive transcranial brain stimulation with neuroimaging and electrophysiology: Current approaches and future perspectives. NeuroImage. 2016 Oct;140:4–19.

10. Polanía R, Nitsche MA, Ruff CC. Studying and modifying brain function with non-invasive brain stimulation. Nat Neurosci [Internet]. 2018; Available from: http://www.nature.com/articles/s41593-017-0054-4

11. To WT, De Ridder D, Hart Jr. J, Vanneste S. Changing Brain Networks Through Non-invasive Neuromodulation. Front Hum Neurosci. 2018 Apr 13;12:128.

12. Weinrich CA, Brittain JS, Nowak M, Salimi-Khorshidi R, Brown P, Stagg CJ. Modulation of Long-Range Connectivity Patterns via Frequency-Specific Stimulation of Human Cortex. Curr Biol. 2017;27(19):3061-3068.e3.

13. Bradley C, Nydam AS, Dux PE, Mattingley JB. State-dependent effects of neural stimulation on brain function and cognition. Nat Rev Neurosci. 2022 Aug;23(8):459–75.

14. Fiene M, Schwab BC, Misselhorn J, Herrmann CS, Schneider TR, Engel AK. Phase-specific manipulation of rhythmic brain activity by transcranial alternating current stimulation. Brain Stimulat. 2020;13(5):1254–62.

15. Fiene M, Radecke JO, Misselhorn J, Sengelmann M, Herrmann CS, Schneider TR, et al. tACS phase-specifically biases brightness perception of flickering light. Brain Stimulat. 2022 Jan;15(1):244– 53.

16. Antal A, Bikson M, Datta A, Lafon B, Dechent P, Parra LC, et al. Imaging artifacts induced by electrical stimulation during conventional fMRI of the brain. NeuroImage. 2014 Jan;85:1040–7.

17. Wischnewski M, Alekseichuk I, Opitz A. Neurocognitive, physiological, and biophysical effects of transcranial alternating current stimulation. Trends Cogn Sci. 2022 Dec;S1364661322002984.

18. Krause MR, Vieira PG, Thivierge JP, Pack CC. Brain stimulation competes with ongoing oscillations for control of spike timing in the primate brain. Luo H, editor. PLOS Biol. 2022 May 25;20(5):e3001650.

19. Liu A, Vöröslakos M, Kronberg G, Henin S, Krause MR, Huang Y, et al. Immediate neurophysiological effects of transcranial electrical stimulation. Nat Commun. 2018 Dec 1;9(1).

20. Kang N, Ko DK, Cauraugh JH. Bimanual motor impairments in older adults: An updated systematic review and meta-analysis. EXCLI J 21Doc1068 ISSN 1611-2156 [Internet]. 2022 [cited 2024 Jul 17]; Available from: https://www.excli.de/index.php/excli/article/view/5236

21. Kim RK, Kang N. Bimanual Coordination Functions between Paretic and Nonparetic Arms: A Systematic Review and Meta-analysis. J Stroke Cerebrovasc Dis. 2020 Feb;29(2):104544.

22. Nettersheim FS, Loehrer PA, Weber I, Jung F, Dembek TA, Pelzer EA, et al. Dopamine substitution alters effective connectivity of cortical prefrontal, premotor, and motor regions during complex bimanual finger movements in Parkinson’s disease. NeuroImage. 2019 Apr;190:118–32.

23. Patel P, Lodha N. Functional implications of impaired bimanual force coordination in chronic stroke. Neurosci Lett. 2020 Nov;738:135387.

24. Schwab BC, Misselhorn J, Engel AK. Modulation of large-scale cortical coupling by transcranial alternating current stimulation. Brain Stimulat. 2019;12(5):1187–96.

25. Brainard DH. The Psychophysics Toolbox. Spat Vis. 1997;10(4):433–6.

26. Bates D, Mächler M, Bolker B, Walker S. Fitting Linear Mixed-Effects Models Using lme4. J Stat Softw [Internet]. 2015 [cited 2024 Jul 30];67(1). Available from: http://www.jstatsoft.org/v67/i01/

27. Thielscher A, Antunes A, Saturnino GB. Field modeling for transcranial magnetic stimulation: A useful tool to understand the physiological effects of TMS? In: 2015 37th Annual International Conference of the IEEE Engineering in Medicine and Biology Society (EMBC) [Internet]. Milan: IEEE; 2015 [cited 2024 Jul 30]. p. 222–5. Available from: http://ieeexplore.ieee.org/document/7318340/

28. Puonti O, Van Leemput K, Saturnino GB, Siebner HR, Madsen KH, Thielscher A. Accurate and robust whole-head segmentation from magnetic resonance images for individualized head modeling. NeuroImage. 2020 Oct;219:117044.

29. Yushkevich PA, Piven J, Hazlett HC, Smith RG, Ho S, Gee JC, et al. User-guided 3D active contour segmentation of anatomical structures: Significantly improved efficiency and reliability. NeuroImage. 2006 Jul;31(3):1116–28.

30. Glasser MF, Coalson TS, Robinson EC, Hacker CD, Harwell J, Yacoub E, et al. A multi-modal parcellation of human cerebral cortex. Nature. 2016 Aug;536(7615):171–8.

31. Tzourio-Mazoyer N, Landeau B, Papathanassiou D, Crivello F, Etard O, Delcroix N, et al. Automated Anatomical Labeling of Activations in SPM Using a Macroscopic Anatomical Parcellation of the MNI MRI Single-Subject Brain. NeuroImage. 2002 Jan;15(1):273–89.

32. Moeller S, Yacoub E, Olman CA, Auerbach E, Strupp J, Harel N, et al. Multiband multislice GE-EPI at 7 tesla, with 16-fold acceleration using partial parallel imaging with application to high spatial and temporal whole-brain fMRI. Magn Reson Med. 2010 May;63(5):1144–53.

33. Feinberg DA, Moeller S, Smith SM, Auerbach E, Ramanna S, Glasser MF, et al. Multiplexed Echo Planar Imaging for Sub-Second Whole Brain FMRI and Fast Diffusion Imaging. Valdes-Sosa PA, editor. PLoS ONE. 2010 Dec 20;5(12):e15710.

34. Xu J, Moeller S, Auerbach EJ, Strupp J, Smith SM, Feinberg DA, et al. Evaluation of slice accelerations using multiband echo planar imaging at 3T. NeuroImage. 2013 Dec;83:991–1001.

35. Jenkinson M, Beckmann CF, Behrens TEJ, Woolrich MW, Smith SM. Fsl. NeuroImage. 2012;62(2):782–90.

36. Jenkinson M, Bannister P, Brady M, Smith S. Improved Optimization for the Robust and Accurate Linear Registration and Motion Correction of Brain Images. NeuroImage. 2002 Oct;17(2):825–41.

37. Salimi-Khorshidi G, Douaud G, Beckmann CF, Glasser MF, Griffanti L, Smith SM. Automatic denoising of functional MRI data: Combining independent component analysis and hierarchical fusion of classifiers. NeuroImage. 2014 Apr;90:449–68.

38. Beckmann CF, Jenkinson M, Smith SM. General multilevel linear modeling for group analysis in FMRI. NeuroImage. 2003 Oct;20(2):1052–63.

39. Fan L, Li H, Zhuo J, Zhang Y, Wang J, Chen L, et al. The Human Brainnetome Atlas: A New Brain Atlas Based on Connectional Architecture. Cereb Cortex. 2016 Aug;26(8):3508–26.

40. Lebihan B, Mobers L, Daley S, Battle R, Leclercq N, Misic K, et al. Bifocal tACS over the primary sensorimotor cortices increases interhemispheric inhibition and improves bimanual dexterity. Cereb Cortex. 2025 Feb 3;bhaf011.

41. Schwab BC, König P, Engel AK. Spike-timing-dependent plasticity can account for connectivity aftereffects of dual-site transcranial alternating current stimulation. NeuroImage. 2021;237(March):118179.

42. Heise KF, Albouy G, Dolfen N, Peeters R, Mantini D, Swinnen SP. Induced zero-phase synchronization as a potential neural code for optimized visuomotor integration. Brain Stimulat. 2025 Mar;S1935861X25000762.

43. Jenkins IH, Jahanshahi M, Jueptner M, Passingham RE, Brooks DJ. Self-initiated versus externally triggered movements. II. The effect of movement predictability on regional cerebral blood flow. Brain. 2000;123(6):1216–28.

44. Lu MK, Arai N, Tsai CH, Ziemann U. Movement related cortical potentials of cued versus self-initiated movements: Double dissociated modulation by dorsal premotor cortex versus supplementary motor area rTMS. Hum Brain Mapp. 2012;33(4):824–39.

45. Mushiake H, Inase M, Tanji J. Neuronal activity in the primate premotor, supplementary, and precentral motor cortex during visually guided and internally determined sequential movements. J Neurophysiol. 1991;66(3):705–18.

46. Callaert DV, Vercauteren K, Peeters R, Tam F, Graham S, Swinnen SP, et al. Hemispheric asymmetries of motor versus nonmotor processes during (visuo)motor control. Hum Brain Mapp. 2011 Aug;32(8):1311–29.

47. Heise KF, Monteiro TS, Leunissen I, Mantini D, Swinnen SP. Distinct online and offline effects of alpha and beta transcranial alternating current stimulation (tACS) on continuous bimanual performance and task-set switching. Sci Rep. 2019 Dec 1;9(1).

48. Lafleur LP, Klees-Themens G, Chouinard-Leclaire C, Larochelle-Brunet F, Tremblay S, Lepage JF, et al. Neurophysiological aftereffects of 10 Hz and 20 Hz transcranial alternating current stimulation over bilateral sensorimotor cortex. Brain Res. 2020 Jan;1727:146542.

49. Rostami M, Lee A, Frazer AK, Akalu Y, Siddique U, Pearce AJ, et al. Determining the effects of transcranial alternating current stimulation on corticomotor excitability and motor performance: A sham-controlled comparison of four frequencies. Neuroscience. 2025 Mar;568:12–26.

50. Schoenfeld MJ, Grigoras IF, Stagg CJ, Zich C. Investigating Different Levels of Bimanual Interaction With a Novel Motor Learning Task: A Behavioural and Transcranial Alternating Current Stimulation Study. Front Hum Neurosci. 2021;15(November):1–16.

51. Hiromitsu K, Asai T, Kadota H, Imaizumi S, Kamata M, Imamizu H. Immediate Modulation of the Blood Oxygenation Level-Dependent Signals by Dual-Site Transcranial Alternating Current Stimulation Propagates Across the Whole Brain [Internet]. 2024 [cited 2024 Sep 13]. Available from: http://biorxiv.org/lookup/doi/10.1101/2024.09.03.610912

52. Vosskuhl J, Huster RJ, Herrmann CS. BOLD signal effects of transcranial alternating current stimulation (tACS) in the alpha range: A concurrent tACS–fMRI study. NeuroImage. 2016 Oct;140:118–25.

53. Vieira PG, Krause MR, Laamerad P, Pack CC. Brain stimulation preferentially influences long-range projections in primates [Internet]. 2025 [cited 2025 Mar 4]. Available from: http://biorxiv.org/lookup/doi/10.1101/2025.02.19.639189

