## Supplementary Information for "Dual-site beta transcranial alternating current stimulation during a bimanual coordination task modulates functional connectivity between motor areas"

### 1 Supplementary Methods

#### 1.1 Magnetic Resonance Imaging

##### 1.1.1 Acquisition

Due to the repositioning of one participant between the fMRI runs, we acquired two field maps and no T2 scan for this participant.

##### 1.1.2 Task-related MR analyses

For the activation analyses, we examined the combination of the following contrasts on the first level: slow+fast and slow-fast; and on the second level: SH-JP, JP-SH, SH-IP, IP-SH, JP-IP, IP-JP for significant clusters. (SH: sham, in-phase: IP, jittered-phase: JP)

For the functional connectivity (FC) analyses, the speed condition contrasts (slow+fast and slow-fast) were modelled in the psychological factor of the psychophysiological interaction analysis (PPI). We examined the combination of the following contrasts on the first level: +PPI (positive FC) and -PPI (negative FC); and on the second level: SH-JP, JP-SH, SH-IP, IP-SH, JP-IP, IP-JP.

Only contrasts mentioned above with significant clusters were reported in tables in the main manuscript. Additionally, for the extraction of the GLM betas for the figures we also ran the contrasts for the individual conditions (i.e., slow/fast and IP/JP/SH).

### 2 Supplementary Results

#### 2.1 Moderate side effects of tACS

The more extensive questionnaire on sensations of tACS that volunteers filled in at the end of the study showed that on a range from 1 to 5 (i.e., 1=absent, 2=mild, 3=moderate, 4=marked, 5=strong), volunteers rated the sensations mild to moderate, as described in Supplementary Table S4. No volunteer described an increase of any sensation over time.

For the sensation ratings after each concurrent tACS-task run on visit 2, we found a significant effect of stimulation in a Kruskal-Wallis test ($X^2(2)=22.032$, $p<0.0001$). Specifically, volunteers rated the sensations as stronger for active tACS (IP and JP) than for the sham stimulation (see Supplementary Figure S6; Bonferroni-corrected post-hoc Dunn's tests: IP vs SH: Z=3.478, $p=0.008$; JP vs SH: Z=4.456, $p<0.0001$; JP vs IP: Z=-0.910239, $p=0.544$). However, this stimulation effect in sensation ratings cannot explain the below-described effects in FC (see the main text for control analyses on the fMRI data).

### 3 Supplementary Tables

*Table S1.* General activation network elicited by the task independent of tACS. Clusters were determined with cluster-forming thresholds of Z = 2.3 and P = 0.01

| Area | Cluster size (voxel) | Maximum *Z*-score | MNI coordinates (mm) of maximum *Z*-statistic | | |
|---|---|---|---|---|---|
| | | | x | y | z |
| ***Slow+fast*** | | | | | |
| Bilateral M1, S1, premotor areas, SMA, frontal medial cortex, frontal orbital cortex, insular cortex, thalamus, striatum, cerebellum | 40376 | 10.6 | 38 | -22 | 54 |
| Left cerebellum | 454 | 5.38 | -34 | -90 | -30 |
| ***Slow>fast*** | | | | | |
| bilateral precentral and postcentral regions, superior parietal regions, SMA, caudate, thalamus, putamen, central opercular cortex, pallidum, insula, cerebellum | 32408 | 8.9 | 38 | -20 | 52 |
| ***Fast>slow*** | | | | | |
| Occipital cortex, precuneus, inferior temporal areas, hippocampus, parahippocampus, amygdala, insula, left cerebellum | 20392 | 5.76 | 28 | -84 | 50 |
| Frontal areas, anterior cingulate cortex | 3827 | 4.85 | 6 | 36 | 22 |

*Table S2.* Task-related fMRI activation independent of speed and tACS condition with cluster-forming thresholds of Z = 2.3 and P = 0.01 within a pre-threshold mask of the upper limb motor regions from the Brainnetome atlas (A4ul). MNI – Montreal Neurological Institute

| Area | Cluster size (voxel) | Maximum *Z*-score | MNI coordinates (mm) of the centre of gravity *Z*-statistic | | |
|---|---|---|---|---|---|
| | | | x | y | z |
| Left M1 | 465 | 9.24 | -31.4 | -23.7 | 61.4 |
| Right M1 | 625 | 10.6 | 35.3 | -18.4 | 58.3 |

*Table S3.* Functional connectivity independent of speed and tACS condition with cluster-forming thresholds of Z = 2.3 and P = 0.01 within the cortical motor regions (precentral and postcentral gyrus, premotor cortices (PM), supplementary motor area (SMA)). MNI – Montreal Neurological Institute

| Area | Cluster size (voxel) | Maximum *Z*-score | MNI coordinates (mm) of maximum *Z*-statistic | | |
|---|---|---|---|---|---|
| | | | x | y | z |
| **1. PPI seeded in the right M1: slow+fast** | | | | | |
| *Positive connectivity (+PPI)* | | | | | |
| Right M1, SMA | 904 | 5.91 | 30 | -20 | 62 |
| *Negative connectivity (-PPI)* | | | | | |
| Left M1, left postcentral gyrus, left PM | 3769 | 6.61 | -42 | -24 | 60 |
| Right M1, right postcentral gyrus | 1015 | 5.89 | 44 | -20 | 52 |
| **2. PPI seeded in the right M1: slow-fast** | | | | | |
| *Negative connectivity (-PPI)* | | | | | |
| Left M1 | 293 | 3.23 | -42 | -20 | 52 |
| **3. PPI seeded in the left M1: slow+fast** | | | | | |
| *Positive connectivity (+PPI)* | | | | | |
| Left M1 | 496 | 4.7 | -32 | -18 | 54 |
| *Negative connectivity (-PPI)* | | | | | |
| Right M1, right postcentral gyrus | 4000 | 7.29 | -42 | -14 | 52 |
| Left M1, left postcentral gyrus | 1010 | 5.65 | -40 | -26 | 56 |
| **4. PPI seeded in the left M1: slow-fast** | | | | | |
| *Negative connectivity (-PPI)* | | | | | |
| Right M1 | 2502 | 4.42 | 28 | -32 | 64 |
| SMA | 747 | 4.37 | 0 | 24 | 54 |

*Table S4.* Sensation ratings at the end of the study. For each sensation, the mean and standard deviation (SD) are shown from a rating in the range from 1 to 5 (i.e., 1=absent, 2=mild, 3=moderate, 4=marked, 5=strong). Additionally, if participants experienced the sensation, they were asked to rate the time course of the sensations (i.e., whether the sensation increased, decreased of stayed constant over time). The % increasing/decreasing/constant refers to the percentage of participants rating the sensation as increasing/decreasing/constant over the course of the stimulation.

| **sensation** | **mean ± SD** | **% increasing sensation** | **% decreasing sensation** | **% constant sensation** |
|---|---|---|---|---|
| itching | 1.88 ± 0.72 | 0 | 57.14 | 42.86 |
| warmth | 2.06 ± 1.29 | 0 | 43.75 | 56.25 |
| sting | 2.56 ± 0.96 | 0 | 68.75 | 31.25 |
| pulsating | 2.56 ± 1.15 | 0 | 50 | 50 |
| pain | 2.09 ± 0.82 | 0 | 68.75 | 31.25 |
| visual | 2.18 ± 0.75 | 0 | 43.75 | 56.25 |

### 4 Supplementary Figures

#### Questionnaire on sensations during tACS

Participant ID:

Date:

Transcranial alternating current stimulation (tACS) is perceived differently for every person. In this questionnaire, we ask you to describe sensations that you had during the stimulation. Please note that we are interested in your **subjective feelings.**

1) Please rate the **strength** of the following **tactile sensations**:

| | | ***absent*** | ***mild*** | ***moderate*** | ***marked*** | ***strong*** |
|---|---|---|---|---|---|---|
| **1** | **Itching** | **O** | **O** | **O** | **O** | **O** |
| **2** | **Warmth** | **O** | **O** | **O** | **O** | **O** |
| **3** | **Sting** | **O** | **O** | **O** | **O** | **O** |
| **4** | **Pulsating** | **O** | **O** | **O** | **O** | **O** |
| **5** | **Pain** | **O** | **O** | **O** | **O** | **O** |

2) During stimulation, your **visual sensations** may be affected. This may include a flickering light or even light flashes. Please rate the **strength** of any visual sensation you may have experienced:

| ***absent*** | ***mild*** | ***moderate*** | ***marked*** | ***strong*** |
|---|---|---|---|---|
| **O** | **O** | **O** | **O** | **O** |

3) Please rate the **time course** of all sensations:

| | | ***increasing*** | ***decreasing*** | ***constant*** |
|---|---|---|---|---|
| **1** | **Itching** | **O** | **O** | **O** |
| **2** | **Warmth** | **O** | **O** | **O** |
| **3** | **Sting** | **O** | **O** | **O** |
| **4** | **Pulsating** | **O** | **O** | **O** |
| **5** | **Pain** | **O** | **O** | **O** |
| **6** | **Visual** | **O** | **O** | **O** |

1

Questionnaire - Date and Version No: 19/06/2020
V1.0

*Figure S1.* Extensive tACS questionnaire filled in at the end of the study. tACS – transcranial alternating current stimulation

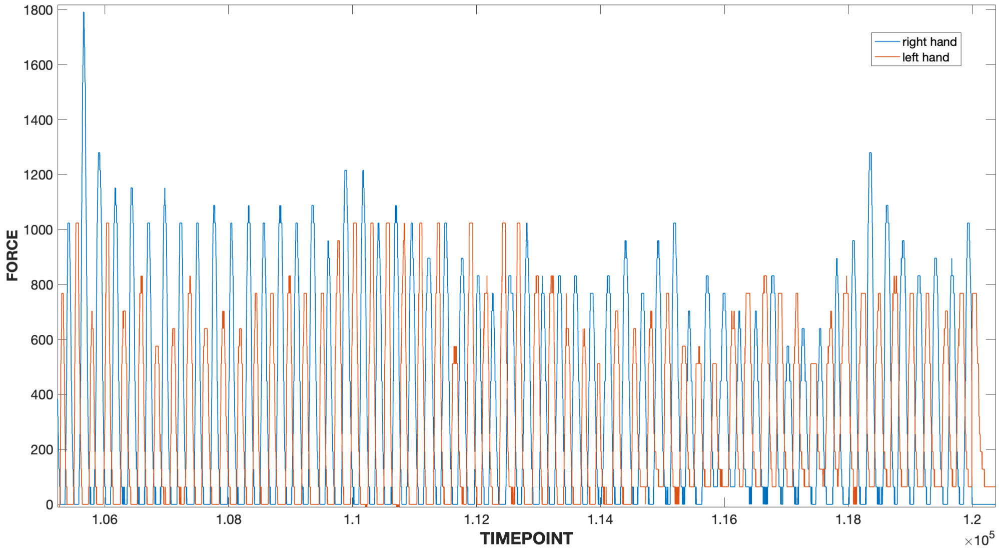


*Figure S2.* Example of the force timecourse of the left and right hand over one task block in one participant.

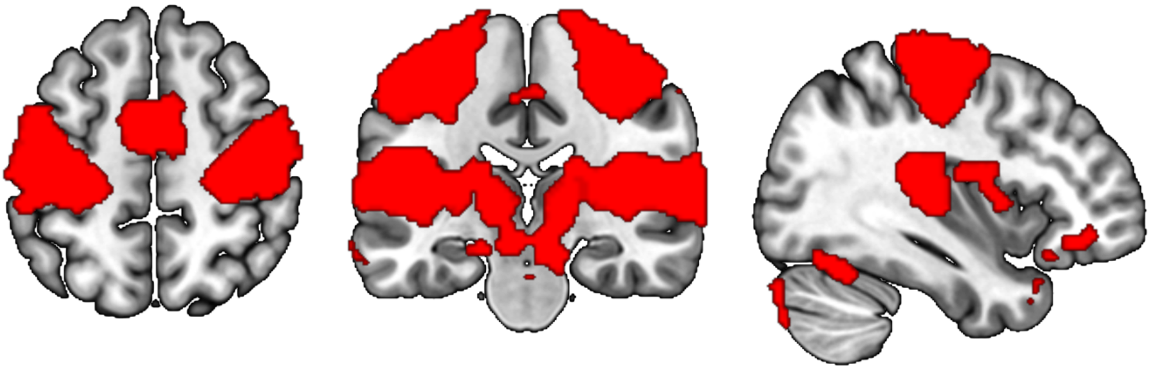

*Figure S3.* General activation network across the whole brain, corresponding to Supplementary Table S1 (slow+fast). The clusters are displayed on a T1-weighted template image.

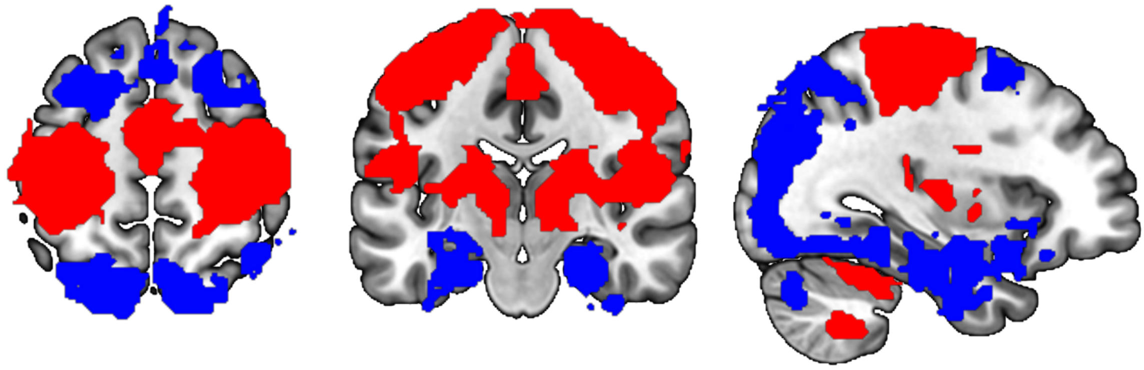

*Figure S4.* Differences in task level activation independent of tACS. Brain regions in red depict higher activation in the slow as compared to the fast conditions (slow>fast) and blue regions depict higher activation for fast as compared to slow conditions (fast>slow) corresponding to Supplementary Table S1. The clusters are displayed on a T1-weighted template image.

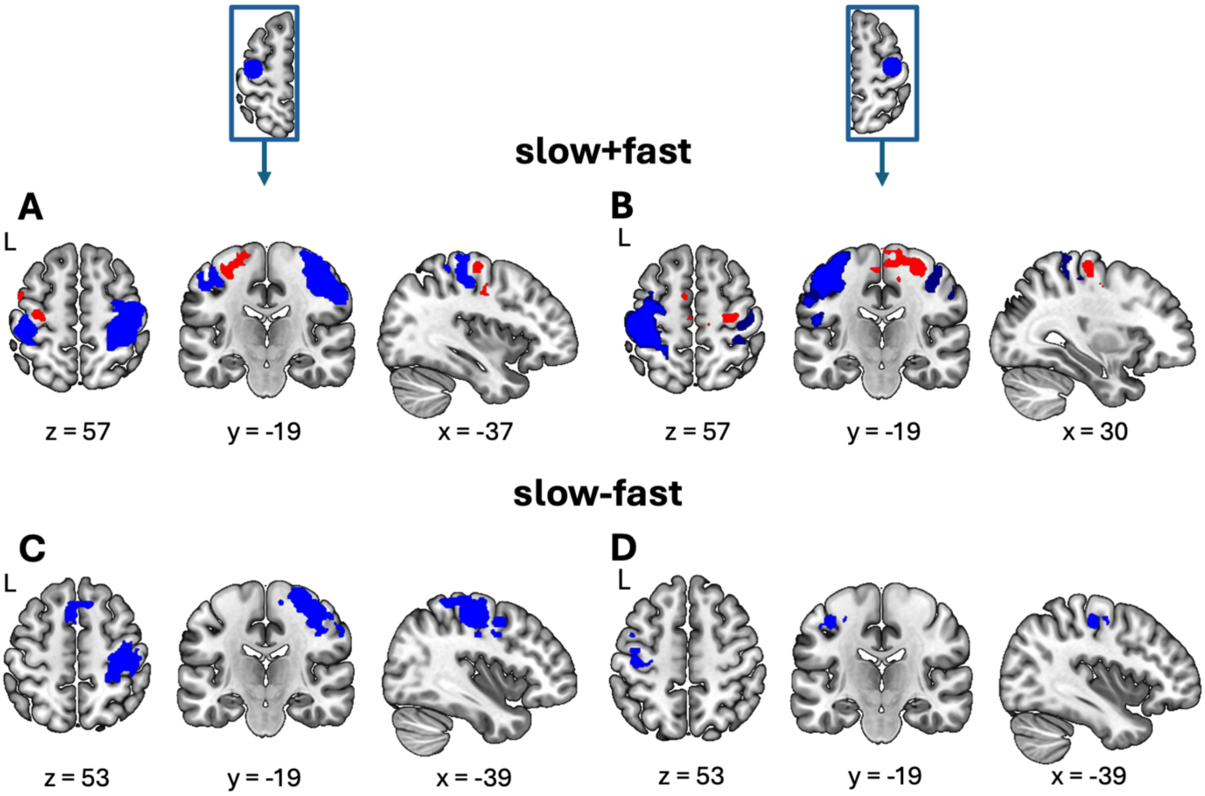


*Figure S5.* Functional connectivity (FC) of the M1 seeds within the cortical motor regions, corresponding to Supplementary Table S3. FC of the left M1 seed is depicted in (A) across speed conditions and in (C) comparing the slow and fast speed conditions. FC of the right M1 seed is depicted in (B) across speed conditions and in (D) comparing the slow and fast speed conditions. Red clusters represent positive FC (+PPI) and blue clusters negative FC (-PPI). The clusters are displayed on a T1-weighted template image. FC – functional connectivity, PPI – psychophysiological interaction

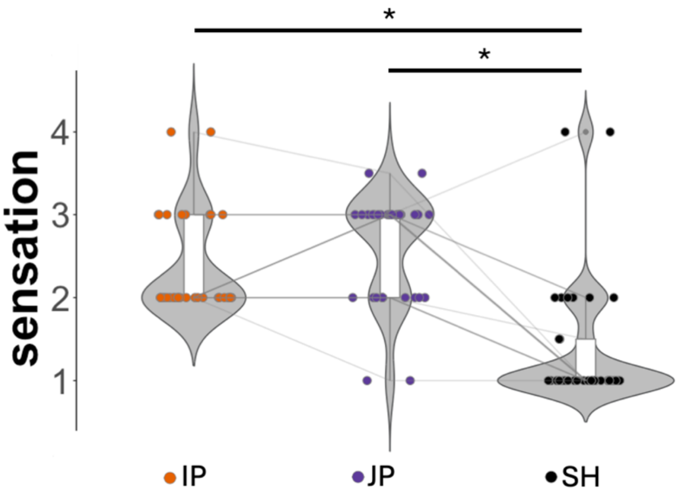


*Figure S6.* Sensations were rated higher for active as compared to the sham tACS conditions. Filled circles represent individual data and lines connect these across conditions for each participant. The shape of the violin plots and boxplots depict the distribution of the data. Asterisks (*) indicate significant statistical tests ($p$<0.05). tACS – transcranial alternating current stimulation, IP – in-phase tACS, JP – jittered-phase tACS, SH – sham tACS
